# A computational model predicts sex-specific responses to calcium channel blockers in mammalian mesenteric vascular smooth muscle

**DOI:** 10.1101/2023.06.24.546394

**Authors:** Gonzalo Hernandez-Hernandez, Samantha C. O’Dwyer, Collin Matsumoto, Mindy Tieu, Zhihui Fong, Pei-Chi Yang, Timothy J. Lewis, L. Fernando Santana, Colleen E. Clancy

## Abstract

The function of the smooth muscle cells lining the walls of mammalian systemic arteries and arterioles is to regulate the diameter of the vessels to control blood flow and blood pressure. Here, we describe an *in-silico* model, which we call the “Hernandez-Hernandez model”, of electrical and Ca^2+^ signaling in arterial myocytes based on new experimental data indicating sex-specific differences in male and female arterial myocytes from murine resistance arteries. The model suggests the fundamental ionic mechanisms underlying membrane potential and intracellular Ca^2+^ signaling during the development of myogenic tone in arterial blood vessels. Although experimental data suggest that K_V_1.5 channel currents have similar amplitudes, kinetics, and voltage dependencies in male and female myocytes, simulations suggest that the K_V_1.5 current is the dominant current regulating membrane potential in male myocytes. In female cells, which have larger K_V_2.1 channel expression and longer time constants for activation than male myocytes, predictions from simulated female myocytes suggest that K_V_2.1 plays a primary role in the control of membrane potential. Over the physiological range of membrane potentials, the gating of a small number of voltage-gated K^+^ channels and L-type Ca^2+^ channels are predicted to drive sex-specific differences in intracellular Ca^2+^ and excitability. We also show that in an idealized computational model of a vessel, female arterial smooth muscle exhibits heightened sensitivity to commonly used Ca^2+^ channel blockers compared to male. In summary, we present a new model framework to investigate the potential sex-specific impact of anti-hypertensive drugs.

## Introduction

Our primary objective was to develop and implement a novel computational model that comprehensively describes the essential mechanisms underlying electrical activity and Ca^2+^ dynamics in arterial myocytes. We aimed to uncover the key components necessary and sufficient to fully understand the behavior of arterial vascular smooth muscle myocytes and the cellular response to variations in pressure. The model represents the first-ever integration of sex-specific variations in voltage-gated K_V_2.1 and Ca_V_1.2 channels, enabling the prediction of sex-specific disparities in membrane potential and the regulation of Ca^2+^ signaling in smooth muscle cells from systemic arteries. To further investigate sex-specific responses to antihypertensive medications, we extended our investigation to include a one-dimensional (1D) representation of tissue. This approach enabled us to simulate and forecast the effects of Ca^2+^ channel blockers within the controlled environment of an idealized mesenteric vessel. It is worth noting that this computational framework can be expanded to predict the consequences of antihypertensive drugs and other perturbations, transitioning seamlessly from single-cell to tissue-level simulations.

Previous mathematical models^1–4^ of vascular smooth muscle myocytes generated to describe the membrane potential and Ca^2+^ signaling in vascular smooth muscle cells have described the activation of G-protein-coupled receptors (GPCRs) by endogenous or pharmacological vasoactive agents activating inositol 1,4,5-trisphosphate (IP_3_) and ryanodine (RyR) receptors resulting in the initiation of calcium waves. Earlier models have also provided insights into the contraction activation by agonists and the behavior of vasomotion. In a major step forward, the Karlin model^5^ incorporated new cell structure data and electrophysiology experimental data in a computational model that predicted the essential behavior of membrane potential and Ca^2+^ signaling arising from intracellular domains found in arterial myocytes. One notable limitation of earlier models is that they are based entirely on data from male animals. Furthermore, many data used to parameterize the Karlin model were obtained from smooth muscle from cerebral arteries. While cerebral arteries are important for brain blood flow, they do not control systemic blood pressure. Furthermore, they do not take into consideration the role of K_V_2.1 channels in the regulation of smooth muscle cell membrane potential.

The function of the smooth muscle cells that wrap around small arteries is to regulate the diameter of these vessels. Arterial myocytes contract in response to increases in intravascular pressure^6^. Based on work largely done using cerebral arterial smooth muscle, a model has been proposed in which this myogenic response is initiated when membrane stretch activates Na^+^-permeable canonical TRPC6^7,8^ and melastatin-type TRPM4^9,10^. A recent study in smooth muscle from mesenteric arteries identified two additional TRP channels to the chain of events that link increases in intravascular pressure to arterial myocyte depolarization: TRPP1 (PKD1) and TRPP2(PKD2) channels^11,12^. Together, these studies point to an elaborate multiprotein complex that plays a critical role in sensing pressure and initiating the myogenic response by inducing membrane depolarization and activating voltage-gated, dihydropyridine-sensitive L-type Ca_V_1.2 Ca^2+^ channels^13,14^. Ca^2+^ entry via a single or small cluster of Ca_V_1.2 channels produces a local increase in intracellular free Ca^2+^ ([Ca^2+^]_i_) called a “Ca_V_1.2 sparklet”^15–18^. Activation of multiple Ca_V_1.2 sparklets produces a global increase in [Ca^2+^]_i_ that activates myosin light chain kinase, which initiates actin-myosin cross-bridge cycling and thus contraction^19^.

Negative feedback regulation of membrane depolarization and Ca^2+^ sparklet activation occurs through the activation of large-conductance, Ca^2+^-activated K^+^ (BK_Ca_) channels as well as voltage-dependent K_V_2.1 and K_V_1.5/1.2 K^+^ channels^20–23^. BK_Ca_ channels are organized into clusters along the sarcolemma of arterial myocytes^24^ and are activated by Ca^2+^ sparks resulting from the simultaneous opening of ryanodine receptors type 2 (RyR2) located in a specialized junctional sarcoplasmic reticulum (SR) ^22,25–28^. Because the input resistance of arterial myocytes is high^29,30^(about 2-10 GΩ), even relatively small currents (10-30 pA) produced by the activation of a small cluster^22,31,32^ of 6-12 BK_Ca_ channels by a Ca^2+^ spark can transiently hyperpolarize the membrane potential of these cells by 10-30 mV. Accordingly, decreases in BK_Ca_, K_V_1.2, K_V_1.5, and/or K_V_2.1 channels depolarize arterial myocytes, increasing Ca_V_1.2 channel activity, [Ca^2+^]_i_, and contraction of arterial smooth muscle^21,33–36^.

A recent study by O’Dwyer *et al.*^20^ suggested that K_V_2.1 channels have dual conducting and structural roles in mesenteric artery smooth muscle with opposing functional consequences. Conductive K_V_2.1 channels oppose vasoconstriction by inducing membrane hyperpolarization. Paradoxically, by promoting the structural clustering of the Ca_V_1.2 channel, K_V_2.1 enhances Ca^2+^ influx and induces vasoconstriction. Interestingly, K_V_2.1 protein is expressed to a larger extent in female than in male arterial smooth muscle. This induced larger Ca_V_1.2 clusters and activity in female than in male arterial myocytes.

Here, we describe a new model, which we call the “Hernandez-Hernandez model”, of mesenteric smooth muscle myocytes that incorporates new electrophysiological and Ca^2+^ signaling data suggesting key sex-specific differences in male and female arterial myocytes. The model simulates membrane currents and their impact on membrane potential as well as local and global [Ca^2+^]_i_ signaling in male and female myocytes. The Hernandez-Hernandez model predicts that K_V_2.1 channels play a critical, unexpectedly large role in the control of membrane potential in female myocytes compared to male myocytes. Importantly, our model predicts that clinically used antihypertensive Ca_V_1.2 channel blockers cause larger reductions in Ca_V_1.2 currents in female than in male arterial myocytes.

Finally, we present a one-dimensional (1D) vessel representation of electrotonically coupled arterial myocytes connected in series. Predictions from the idealized vessel suggest that Ca^2+^ channel blockers are more potent in females resulting in a more substantial [Ca^2+^]_i_ reduction in female arterial smooth muscle compared to male. The Hernandez-Hernandez model demonstrates the importance of sex-specific differences in Ca_V_1.2 and K_V_2.1 channels and suggests the fundamental electrophysiological and Ca^2+^linked mechanisms of the myogenic tone. The model also points to testable hypotheses underlying differential sex-based pathogenesis of hypertension and distinct responses to antihypertensive agents.

## RESULTS

In this study, we developed a computational model of the electrical activity of an isolated vascular smooth muscle cell (**Figure 1**). A key goal was to optimize and validate the model with experimental data and then use the model to predict the effects of measured sex-dependent differences in the electrophysiology of smooth muscle myocytes.

**Figure 1.**
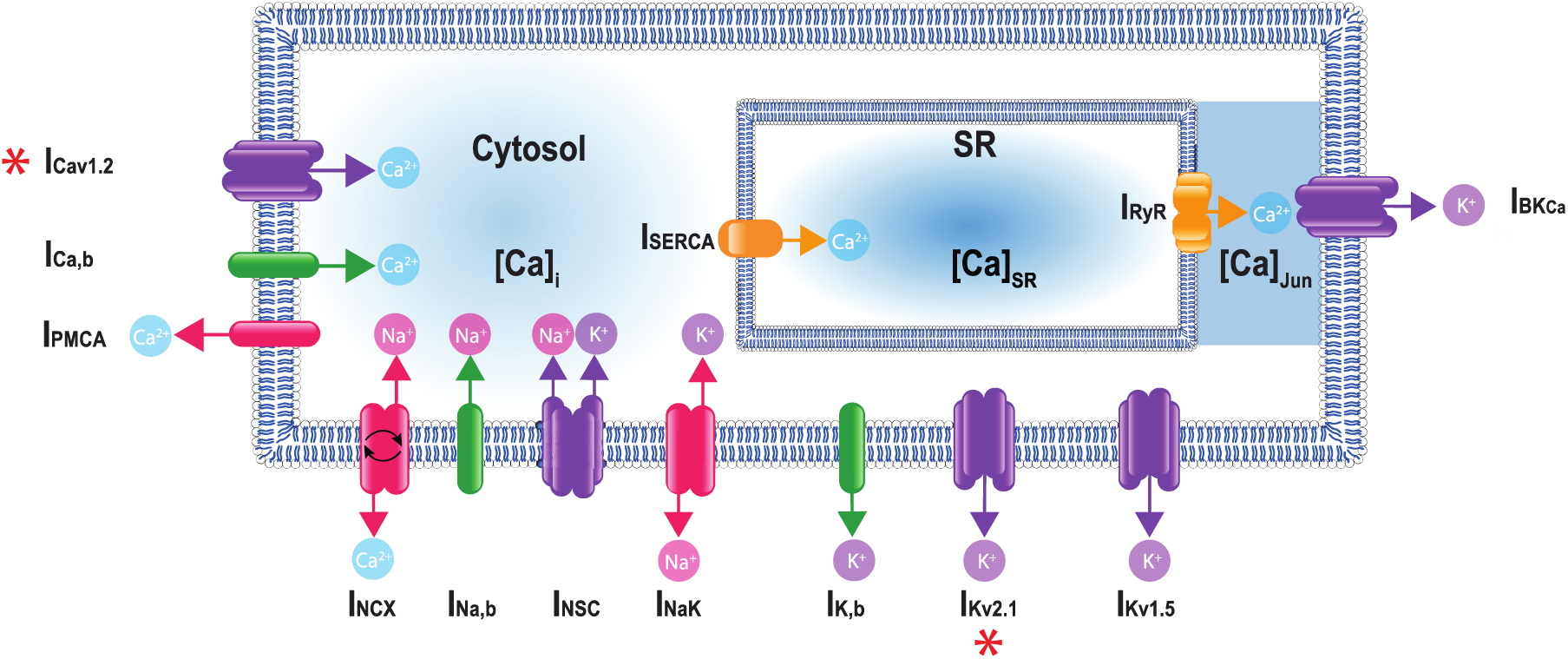
A schematic representation of the Hernandez-Hernandez model. The components of the model include major ion channel currents shown in purple including the voltage-gated L-type calcium current (I_Ca_), nonselective cation current (I_NSC_), voltage-gated potassium currents (I_Kv1.5_ and I_Kv2.1_), and the large-conductance Ca^2+^-sensitive potassium current (I_BKCa_). Currents from pumps and transporters are shown in red including the sodium/potassium pump current (I_NaK_), sodium/calcium exchanger current (I_NCX_), and plasma membrane ATPase current (I_PMCA_). Leak currents are indicated in green including the sodium leak current (I_Na,b_), potassium leak current (I_K,b_), and calcium leak current (I_Ca,b_). In addition, two currents in the sarcoplasmic reticulum are shown in orange: the sarcoplasmic reticulum Ca-ATPase current (I_SERCA_) and ryanodine receptor current (I_RyR_). Calcium compartments comprise three discrete regions including cytosol ([Ca]_i_), sarcoplasmic reticulum ([Ca]_SR_), and the junctional region ([Ca]_Jun_). Red stars (*) indicate measured sex-specific differences in ionic currents.

In constructing the model, we first set out to measure the kinetics of the voltage-gated L-type Ca_V_1.2 currents (I_Ca_) in male and female myocytes using Ca^2+^ as the charge carrier as shown in **Figure 2**. These data provided information on the kinetics of Ca^2+^-dependent activation and inactivation of I_Ca_. I_Ca_ is critical in determining cytosolic concentration [Ca^2+^]_i_ in vascular mesenteric smooth muscle cells and is the predominant pathway for Ca^2+^ entry^13,15,16,18,28,52^. Experiments using whole-cell patch-clamp were undertaken to measure the time constants of activation and deactivation (**panel 2A**) and inactivation (**panel 2B)** in male and female mesenteric artery smooth muscle cells shown as black and blue symbols, respectively. While the data from male (n = 10) and female (n = 12) myocytes showed comparable activation time constants, there was an observable trend of faster inactivation in the female cells in the lower voltage range, but the differences were not statistically significant. Steady-state activation and inactivation were also measured as shown in **panel 2C**, with male data in black symbols and female as blue symbols. No observable differences in the gating characteristics of the male and female I_Ca_ were measured. Finally, the current-voltage relationship is shown from measurements in female (blue) and male (black) in **panel 2D.** This analysis suggests that the amplitude of I_Ca_ was larger in female than in male myocytes over a wide range of membrane potentials.

**Figure 2.**
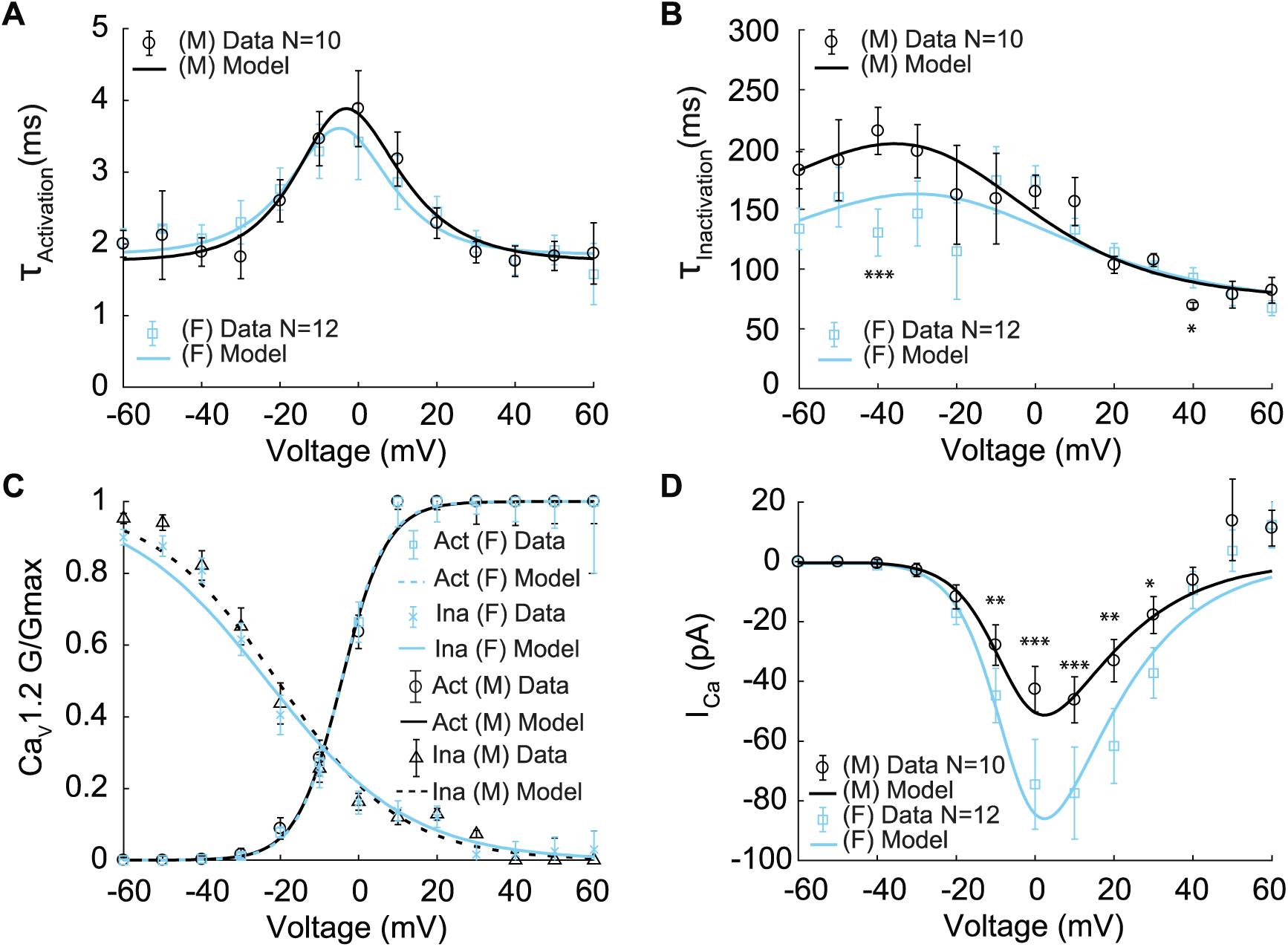
Experimentally measured and modeled L-type calcium currents (I_Ca_) from male and female vascular smooth muscle cells. Properties of I_Ca_ are derived from measurements in male and female vascular smooth muscle (VSM) cells isolated from the mouse mesenteric arteries following voltage-clamp steps from −60 to 60 mV in 10 mV steps from a −80 mV holding potential. Experimental data is shown in black circles for male and blue squares for female. Model fits to experimental data are shown with black solid lines for male and blue solid lines for female. (**A**) Male and female time constants of I_Ca_ activation. (**B**) Male and female time constants of I_Ca_ inactivation. (**C**) Male and female voltage-dependent steady-state activation and inactivation of I_Ca_. (**D**) Current-voltage (I-V) relationship of I_Ca_ from male and female vascular smooth muscle myocytes. *P < 0.05, **P < 0.01, ***P<0.001. Error bars indicated mean ± SEM.

We next used the experimental measurements to build and optimize a Hodgkin-Huxley model based on the data described above. The model includes voltage-dependent activation and inactivation gating variables, dL and dF, respectively. We modeled both gates following the approach by Kernik *et al.* ^45^. It is important to note that smooth muscle cells operate within a voltage regime defined by the window current, which ranges between −45 mV and −20 mV. Under these conditions, [Ca^2+^]_i_ remains below 1 μM. Therefore, we did not consider the Ca^2+^-dependent inactivation gating mode of the channel^2,53^.

The model of I_Ca_ is described by:

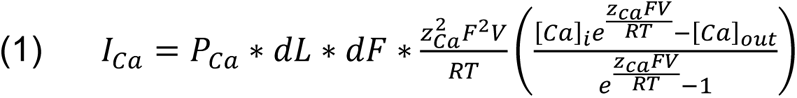

Where P_Ca_ is the ion permeability, R is the gas constant, F is the Faraday’s constant, and z_Ca_ is the valence of the Ca^2+^ ion. Parameters were optimized to male and female experimental data as shown for activation time constants (T_activation_) and inactivation time constants (T_inactivation_) as solid lines in **Figure 2A** and **Figure 2B**, respectively. Model optimization to male and female activation and inactivation curves are shown in **Figure 2C**. The model was also optimized to the I_Ca_ current-voltage (I-V) relationships shown as solid lines in **Figure 2D**.

We next set out to determine sex differences in voltage-gated K^+^ currents (I_K_) in male and female mesenteric smooth muscle cells. I_K_ is produced by the combined activation of K_V_ and BK_Ca_ channels. Following the approach previously published by O’Dwyer *et al.,*^20^ we quantified the contribution of K_V_ (I_KV_) and BK_Ca_ (I_BKCa_) current to I_K_. K^+^ currents were recorded before and after the application of the channel blocker iberiotoxin (IBTX; 100nm). Once identified the contribution I_BKCa_ current, we isolated the voltage-gated potassium currents (I_KV_) whose contributors include the voltage-gated potassium channels K_V_1.5 and K_V_2.1. The presumed function of K_V_1.5 and K_V_2.1 channels on membrane potential is to produce delayed rectifier currents to counterbalance the effect of the inward currents^19,20^.

Having isolated I_KV_, K_V_2.1 currents were identified using the application of the K_V_2.1 blocker ScTx1 (100 nM). As a result, the remaining ScTx1-insensitive component of the I_KV_ current was attributed to K_V_1.5 channels. The results are shown in **Figure 3**. Experiments using whole-cell patch-clamp were undertaken to measure the steady-state activation G/G_max_ of the K_V_2.1current (I_Kv2.1_) as shown in **panel 3A** in female (blue) and male (black) myocytes and no significant differences were observed. Measurements of time constants of activation (**panel 3B**) of I_Kv2.1_ in the voltage range of −30 to +40 mV in female (blue, n=10) and male (black, n=7) myocytes exhibited significant differences. Notably, activation time constants in male myocytes were smaller than those in female myocytes, corresponding to a faster activation rate in males. The current-voltage relationship of I_Kv2.1_ is shown from measurements in female (blue, n=20) and male (black, n=10) myocytes in **panel 3C**. Significant differences were observed in I_Kv2.1_ at various voltages. In **panel 3C**, data points without asterisks are not considered significant. Similarly, we measured the steady-state activation of the K_V_1.5 current (I_Kv1.5_) as shown in **panel 3D** where male and female experimental data in myocytes are shown with blue and black symbols. Properties of I_Kv1.5_ steady-state activation G/G_max_ show minimal sex-specific differences. The current-voltage relationship of I_Kv1.5_ is shown from measurements in female (blue, n=10) and male (black, n=7) myocytes in **panel 3E.** Finally, the current-voltage relationship of the contribution from I_Kv1.5_ and I_Kv2.1_ to the total voltage-gated current (I_KVTOT_) is shown in **panel 3F** with male and female data shown with black and blue symbols, respectively. Data points in **panel D**, **panel E**, and **panel F** without asterisks are not significant. The table in **panel 3H** summarizes the sex-dependent maximal conductance and the current response at specific voltages of −50 mV, −40 mV, −30 mV, and −20 mV for both I_Kv1.5_ and I_Kv2.1_.

**Figure 3.**
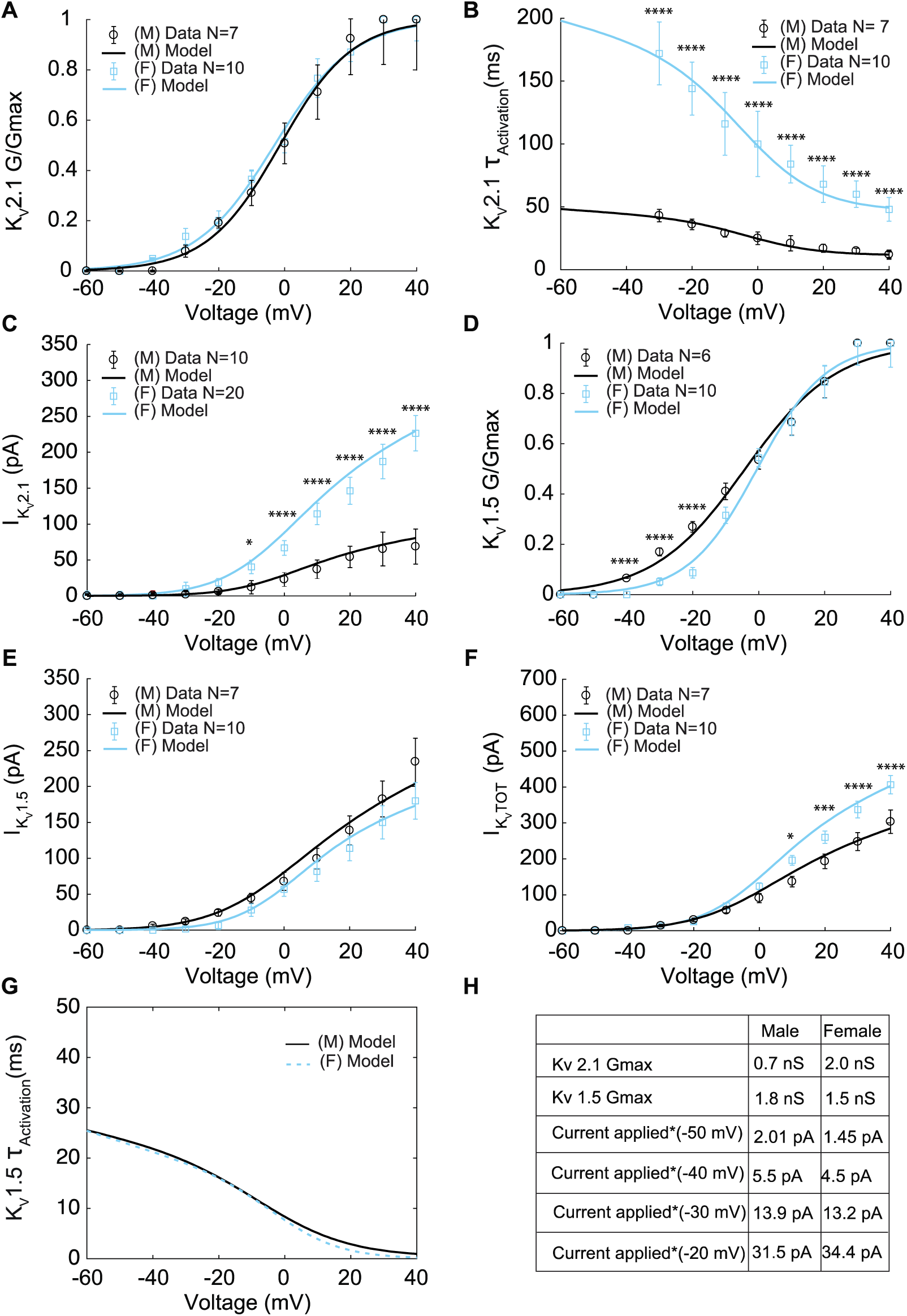
Experimentally measured and modeled potassium currents (I_KvTOT_) from male and female vascular smooth muscle cells. The properties of I_Kv1.5_ and I_Kv2.1_ from experimental measurements in male and female vascular smooth muscle cells isolated from the mouse mesenteric arteries were recorded in response to voltage-clamp from - 60 to 40 mV in 10 mV steps (holding potential −80mV). Experimental data is shown as black circles for male and blue squares for female. Model fits to experimental data are shown with black solid lines for male and blue solid lines for female. (**A**) Male and female voltage-dependent steady-state activation of I_Kv2.1_. (**B**) Male and female time constants of I_Kv2.1_ activation. (**C**) Current-voltage (I-V) relationship of I_Kv2.1_ from male and female myocytes. (**D**) Male and female voltage-dependent steady-state activation of I_Kv1.5_. (**E**) Current-voltage (I-V) relationship of I_Kv1.5_ from male and female myocytes. (**F**) Male and female total voltage-gated potassium current I_KvTOT_ = I_Kv1.5_ + I_Kv2.1_. (**G**) Predicted male and female time constants of the I_Kv1.5_ activation gate. (**H**) Table showing sex-specific differences in conductance and steady-state total potassium current-voltage dependence. *P < 0.05, **P < 0.01, ***P<0.001, ****P<0.0001. Data points without asterisks are not significant. Error bars indicated mean ± SEM.

To understand the contribution of each K^+^ current to the total voltage-gated current (I_KVTOT_) in mesenteric vascular smooth muscle cells we built and optimized a Hodgkin-Huxley model to the data described above. First, we developed a model to describe the K_V_2.1 current. The optimized model to K_V_2.1 experimental data contains only a voltage-dependent activation gating variable (X_2.1act_). Since inactivation time is slow and is well estimated by steady-state^3^, we did not consider its effects in our model. The model of I_Kv2.1_ is described by:

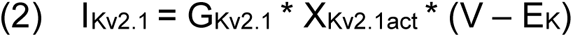

Where *G*__*K*_2.1_ is the maximal conductance of K_V_2.1 channels and E_K_ is the Nernst potential for potassium. Parameters were optimized to male and female experimental data as shown for activation curves in **Figure 3A**. Model optimization to male and female time constants of activation (K_V_2.1 T_Activation_) are shown as solid lines in **Figure 3B**. The model was also optimized to the I_Kv2.1_ current-voltage (I-V) relationships shown as solid lines in **Figure 3C**.

Similarly, we developed a model for K_V_1.5. The model was optimized to the K_V_1.5 experimental data and contains only a voltage-dependent activation gating variable (X_Kv1.5act_). The model of I_Kv1.5_ is described by:

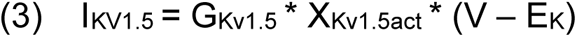

*G*__*K*_1.5_ is the maximal conductance of K_V_1.5 channels and E_K_ is the Nernst potential for potassium. Parameters were optimized to male and female experimental data as shown for activation curves in **Figure 3D**. The model was also optimized to the I_Kv1.5_ current-voltage (I-V) relationships shown as solid lines in **Figure 3E**. From experiments, we optimized the model to reproduce the overall time traces of K_V_ currents. The model predicted that male and female myocytes have comparable time constants of activation in I_Kv1.5_ as shown in **Figure 3G**. Finally, the optimized model of the total voltage-gated current (I_KVTOT_) is shown in **Figure 3F**. The total voltage-gated K^+^ current (I_KvTOT_) is the sum of I_KV1.5_ and I_Kv2.1_ mathematically described as:

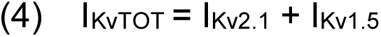

Notably, the main specific sex-specific differences observed in the total voltage-gated K^+^ current (I_KvTOT_) is attributable to the sex-specific differences in the current produced by K_V_2.1 channels.

We next analyzed the contribution of large-conductance calcium-activated potassium (BK_Ca_) channels to vascular smooth muscle cell electrophysiology. BK_Ca_ channels are activated by membrane depolarization or increased [Ca^2+^]_i_ and are expressed in the membrane of vascular smooth muscle cells with α and β1 subunits^22,54,55^. In smooth muscle cells, Ca^2+^ sparks are the physiological activators of BK_Ca_ channels. We relied on the assumption by Tong *et al*.^56^ that BK_Ca_ currents (I_BKCa_) are produced by two current subtypes, one consisting of α subunits (I_BKα_) and the other consisting of α and β1 subunits (I_BKαβ1_). Experimental evidence indicates that BK_Ca_ channels with αβ1 subunits form clusters in the plasma membrane in specialized junctional domains formed by the sarcoplasmic reticulum and the sarcolemma. BK_Ca_ channels with αβ1 subunits colocalize with ryanodine receptors (RyRs) to in the junctional domains. During a Ca^2+^ spark, [Ca^2+^]_i_ elevations ranging from 10 to 100 μM activate BK_Ca_ channels^38,39,52,57,58^. In our model, Ca^2+^ sparks are the physiological activators of BK_Ca_ channels.

The mathematical formulation of the BK_Ca_ with αβ1 current (I_BKαβ1_) was optimized to fit the experimental whole-cell electrophysiological data from Bao and Cox^54^ obtained at room temperature with a BK_Ca_ channel α subunit clone from mSlo-mbr5 and a β1 subunit clone from bovine expressed in *Xenopus laevis* oocytes^54^. Experimental data for steady-state activation and time constants of activation are shown in **Panel 4A** and **Panel 4B** respectively. The activation gating variable (*X*_*ab*_) depends on both voltage and junctional calcium ([Ca^2+^]_Jun_). The activation gate was adapted from the Tong-Taggart model^56^. The model of I_BKαβ1_ is described by:

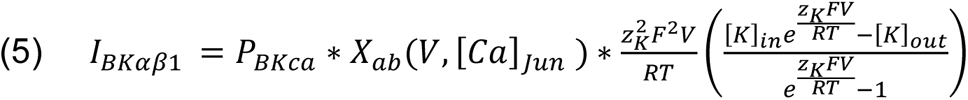

Where P_BKCa_ is the BK_Ca_ ion permeability, R is the gas constant, F is Faraday’s constant, and z_K_ is the valance of the potassium ions. Model optimization to activation curves are shown with solid lines in **Figure 4A** at three different [Ca^2+^]_Jun_ concentrations 1 μM, 10 μM, and 100 μM. The results from the steady-state activation measurements at 10 μM are also in agreement with the experimental data in vascular myocytes in *bufo marinus*^58^ (green symbols) which suggests that BK_Ca_ channels are exposed to a mean junctional Ca^2+^ concentration ([Ca^2+^]_Jun_) of 10 μM. Time constants of activation were measured experimentally at [Ca^2+^]_Jun_ = 0.003 μM, our model was optimized and fit under the same conditions shown in **Figure 4B** as solid lines. Notably when the model was run under predicted [Ca^2+^]_Jun_ = 10 μM conditions as shown in **Figure 4B** dashed lines, there was no effect of the change in [Ca^2+^]_Jun_ on the time constant. The predicted current-voltage (I-V) relationships of I_BKαβ1_ are shown in **Figure 4C** using three different [Ca^2+^]_Jun_ concentrations; 1 μM, 10 μM, and 100 μM. We observed that the I-V curves are similar at [Ca^2+^]_Jun_ concentrations of 10 μM (black trace) and 100 μM (orange trace) but markedly reduced when [Ca^2+^]_Jun_ = 1 μM (blue trace). As expected, the amplitude of the current shown in the I-V curves in **Figure 4D** is sensitively dependent on the number of BK_Ca_ channels as shown, we set [Ca^2+^]_Jun_= 10 μM and simulated the I-V curves using a BK_Ca_ cluster size of 4, 6, 8 and 10 channels.

**Figure 4.**
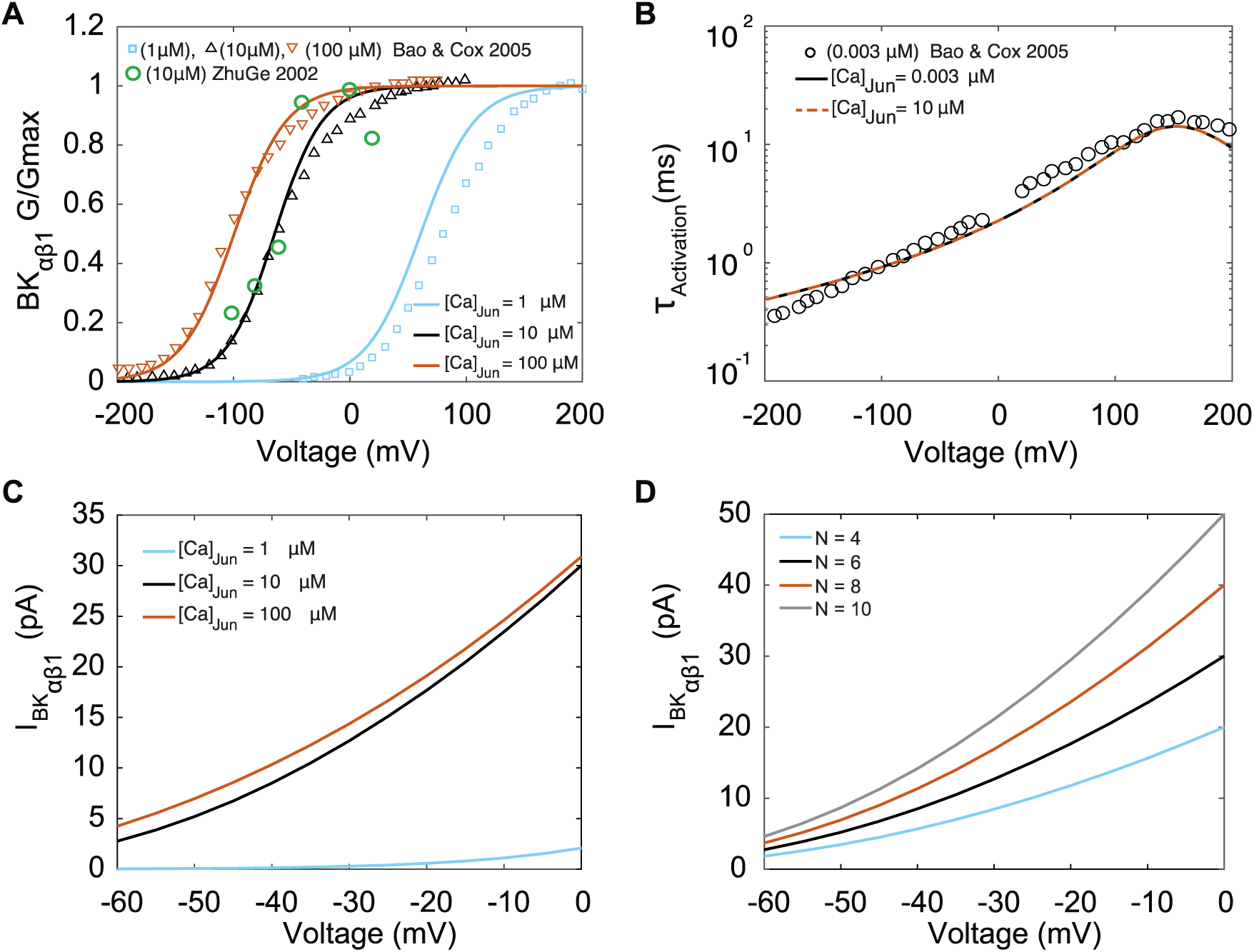
Experimentally measured and modeled large-conductance Ca^2+^-activated K^+^ currents (I_BKαβ1_). The model was optimized to data from Bao and Cox (Bao & Cox, 2005). (**A**) Voltage-dependent activation of I_BKαβ1_ from experiments performed with three different [Ca]_Jun_ concentrations (1 μM, 10 μM, 100 μM) shown in green circles is the data from (Zhuge *et al.*, 2002) (**B**) Voltage-dependent activation time constants with [Ca]_Jun_=0.003 μM and simulations [Ca]_Jun_=10 μM. (**C**) Simulated I-V curve at different peak levels of [Ca]_Jun_ levels. (**D**) Simulated I-V curve with different BK_ca_ average cluster sizes (N = 4,6, 8, and 10).

In vascular smooth muscle cells, the membrane potential over the physiological range of intravascular pressures is less negative than the equilibrium potential of potassium (E_K_ = −84 mV), suggesting active participation of inward currents regulated by sodium conductance^19,59,60^. It has been postulated that basally activating TRP channels generate nonselective cations currents (I_NSC_) that depolarize the membrane potential. We built a model for I_NSC_ as linear and time-independent cation current permeable to K^+^ and Na^+^ with permeability ratios P_Na_: P_K_ =0.9:1.3 adapted from Tong-Taggart model with a reversal potential (E_NSC_) described by:

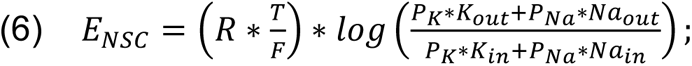

where R is the gas constant, F is the Faraday’s constant, T is the temperature, Na_in_ and K_in_ are the intracellular sodium and potassium intracellular concentrations. Similarly, Na_out_ and Na_out_ are the extracellular sodium and potassium concentrations. The model of I_NSC_ is described by

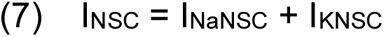

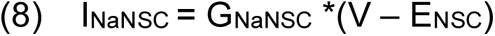

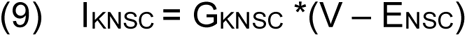

Where I_NaNSC_ represents sodium current contribution, I_KNSC_ represents potassium current contribution, and G_NaNSC_ and G_KNSC_ are the maximal conductances of the contributing sodium and potassium currents. In addition, we also included models for leak currents of ion i calculated as

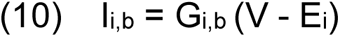

Where the Nernst potential of ion i with valance z_i_ is given by:

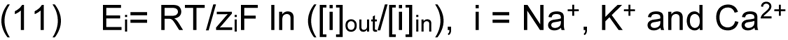

Where R is the gas constant, F is the Faraday’s constant, T is the temperature and [i]_out_ denotes the extracellular concentration of ion i. FF

The remaining ionic currents, pumps, and transporters were optimized to data available in the experimental literature and/or taken from computational models of vascular smooth muscle and cardiac cells. The sodium-potassium pump (I_NaK_) current was modeled using data from smooth muscle cells from mesenteric resistance arteries of the guinea pig^56,61^ and the voltage dependency was adapted from the Luo-Rudy II model^41^. The sodium-calcium exchanger current (I_NCX_) was adapted from the formulation in the ten Tusscher model^62^ and the Luo-Rudy II model^41^. Finally, the sarcolemma calcium pump (I_PMCA_) current was adapted from the Kargacin model^63^.

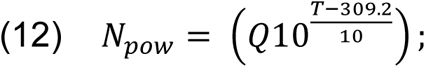

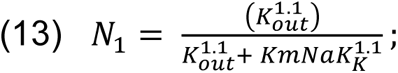

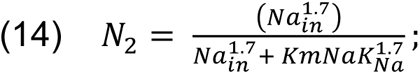

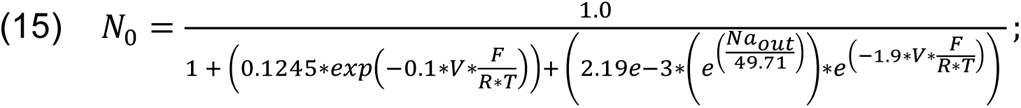

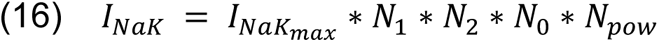

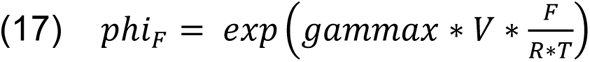

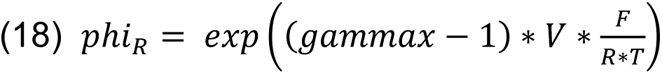

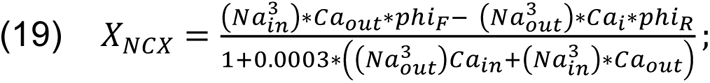

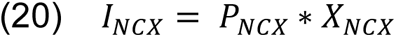

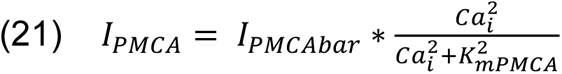

We next set out to connect the ionic models and models of Ca^2+^ handling to make predictions in the whole cell. In **Figure 5**, experimental data indicate that the electrical activity of isolated mesenteric smooth muscle cells in male and female myocytes recorded in current-clamp mode, is characterized by an oscillating membrane potential under physiological conditions. The membrane potential is marked by repetitive spontaneous transient hyperpolarization (TH), a ubiquitous feature of vascular smooth muscle cells^57,64–66^ as shown in **panel 5A.** Both male (black trace) and female (blue trace) myocytes exhibited membrane hyperpolarizing transients in the potential range of −50 to −20 mV. Notably, we observed that female myocytes always maintained a higher depolarizing state between the hyperpolarization events compared to male myocytes.

**Figure 5.**
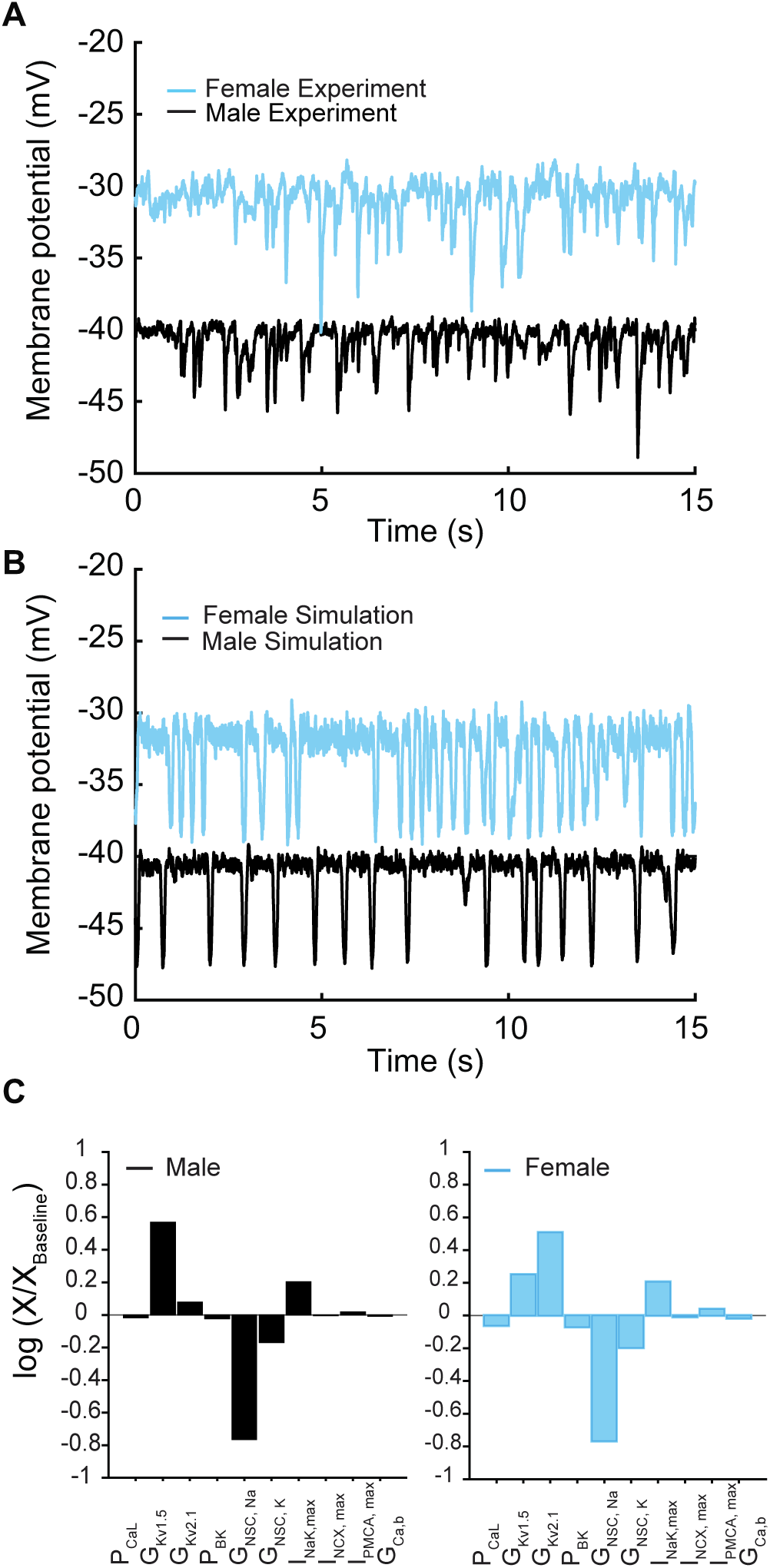
Membrane potential from experiments and simulations in male and female vascular smooth muscle myocytes. **A**) Whole-cell membrane potential recordings in male and female myocytes showing spontaneous repeat transient hyperpolarization of the membrane potential. **B**) Simulated whole-cell membrane potential with physiological noise. **C**) Comparison of sensitivity analysis performed around the baseline membrane potential in male and female models using multivariable regression.

We assessed the predictive capacity of our *in silico* model by comparing it to experimental data. We first compared the morphology of the membrane potential in experiments **panel 5A** versus simulations **panel 5B** in male and female myocytes. Upon comparative analysis between male and female experimental data and simulations, we noted that the baseline membrane potential for male myocytes was around –40 mV, while female myocytes exhibited a slightly more depolarized membrane potential at approximately –30 mV. Despite these variations in baseline membrane potential, both male and female myocytes presented similar peak hyperpolarization values of approximately 10-15 mV, ranging from –50 mV to –30 mV. Similarly, the frequency of THs from multiple myocytes was calculated to be 1 to 2.8 Hz in the range of –50 mV to –30 mV which is identical to the simulated frequency.

In the physiological range in which smooth muscle cells operate (−50 to −20 mV), ionic currents are small and produced by the activation of a small number of ion channels. Local fluctuations in the function of ion channels lead to noisy macroscopic signals that are important to the variability of vascular smooth muscle cells^19^. In addition, smooth muscle cells are subject to high input resistance where small perturbations can lead to large changes in the membrane potential^19,29^. To approximate the physiological realities, we applied two sources of noise to our deterministic *in silico* model to simulate the stochastic fluctuations. The first source of the noise was introduced by adding a fluctuating current term to the differential equations describing changes in membrane potential (dV/dt) which represents the combined effect of the stochastic activity of ion channels in the plasma membrane^46^. Second, we introduced noise into the [Ca]_SR_ to replicate the physiological responses consistent with those observed in experimental studies^40^. Simulated whole-cell membrane potential with physiological noise is shown in **Figure 5B** in male (black trace) and female (blue trace) myocytes.

We conducted a sensitivity analysis to determine which model parameters could underlie the sex-specific differences observed in the experimental data. It is important to note that we have experimental data indicating the amplitude and kinetics for a variety of currents in male and female myocytes. For this reason, those model components were fit to the data, fixed, and were not subject to sensitivity analysis. Our analysis, which focused solely on variations in maximal conductance and maximal ion transport rates of the transmembrane currents, indicated that the non-selective cation currents (I_NSC_) and delayed rectifier currents (I_KvTOT_ = I_Kv2.1_ + I_Kv1.5_) interact to regulate the baseline membrane potential in both male and female vascular smooth muscle myocytes (**Figure 5C**). Given that I_KvTOT_ responds to depolarization, the primary stimulus that triggers depolarization was determined to be attributable solely to the non-selective cation currents (I_NSC_). Indeed, when we adjusted the conductance of the non-selective cation currents and implemented an increase in the conductance of I_NSC_ in the female model, we readily reproduced the sex-specific baseline membrane potential observed experimentally (**Figure 5A)**.

Next, using the whole-cell vascular smooth muscle myocyte computational model, we investigated the sex-specific differences in the contribution to total voltage-gated current (I_KVTOT_) in mesenteric vascular smooth muscle cells. An interesting prediction from the *in silico* simulations is that at different depolarizing states (−45, −40, and −35 mV) induced by changing the conductance of nonselective cationic leak currents (I_NSC_), the contribution of I_Kv2.1_ and I_Kv1.5_ to I_KvTOT_ is different based on sex. In male vascular myocytes, the contribution to total voltage-gated current (I_KVTOT_) is largely attributable to the current produced by K_V_1.5 channels as shown in the lower panel in **Figure 6A**. Our results are consistent with previous studies^35,67,68^ in animal rodent male models showing the characteristic behavior of I_Kv1.5_ to control membrane potential. However, the model predicts that in female myocytes, the contribution to total voltage-gated current (I_KVTOT_) is largely provided by the current produced by K_V_2.1 channels as shown in the upper panel in **Figure 6B**. To illustrate this point quantitatively, at a membrane potential of −40 mV, the contribution of I_KVTOT_ from I_Kv1.5_ and I_Kv2.1_is 86% and 14%, respectively, in male myocytes compared to female myocytes in which the contribution from I_Kv1.5_ and I_Kv2.1_ is 23% and 77%, respectively. Regardless of the depolarization state at −45, −40, or −35 mV, the profiles for male and female myocytes remain essentially the same as shown in **Figures 6C, 6D, and 6E**. The *in silico* simulations suggest a distinctive sex-based function of K_V_1.5 and K_V_2.1 channels that produce the delayed rectifier currents to counterbalance the effect of inward currents causing graded membrane potential depolarizations.

**Figure 6.**
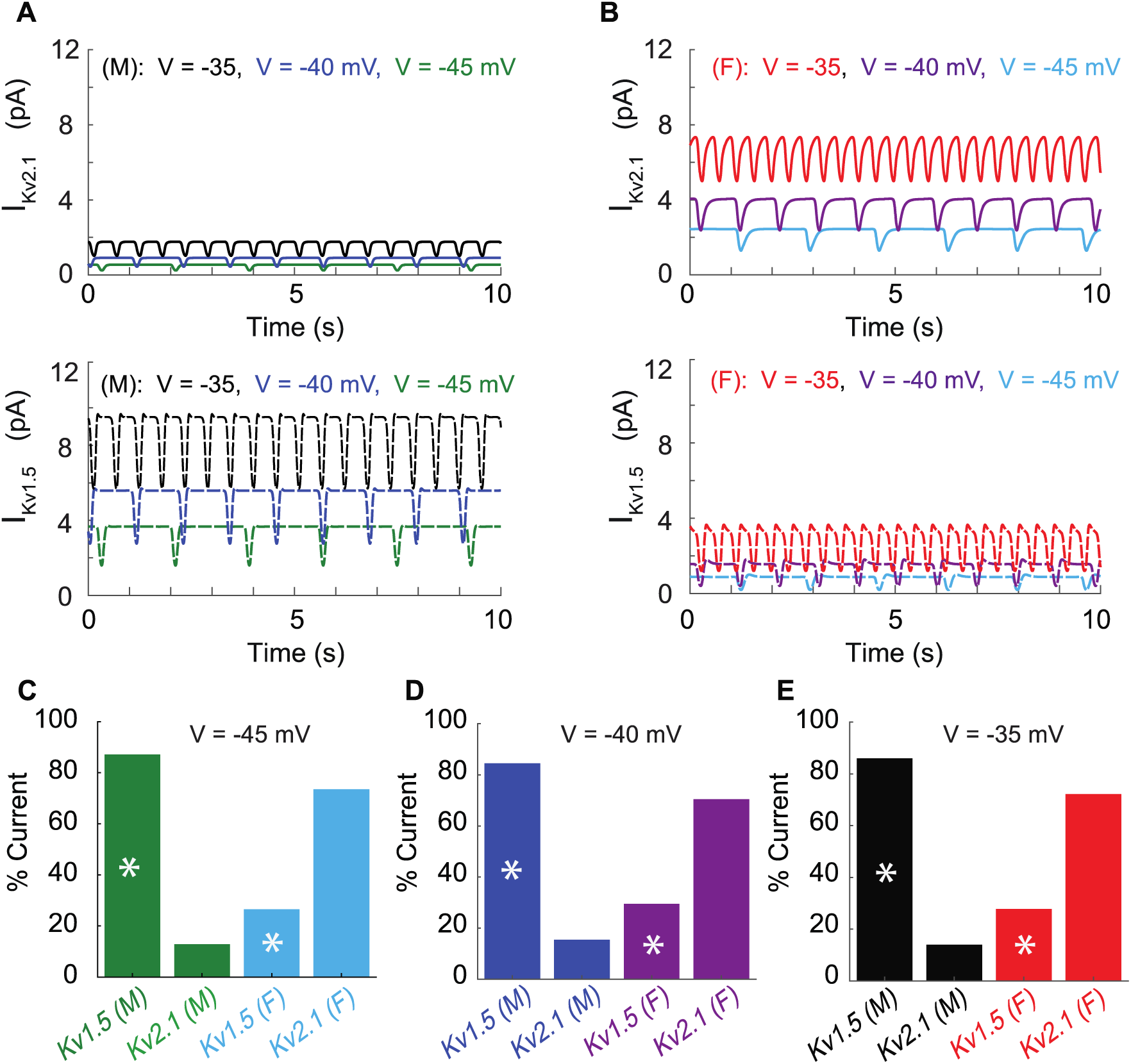
Differential effects of voltage-gated potassium current (I_KVTOT_) block in male and female myocytes. **A**) Simulated time course of male I_Kv2.1_ (*top panel, solid traces*) and I_Kv1.5_ (*lower panel, dashed traces*) at three different baseline membrane potentials (−45 mV *green*, −40 mV *blue*, and −35mV *black*). **B**) Simulated time course of female I_Kv2.1_ (*top panel solid traces*) and I_Kv1.5_ (*lower panel, dashed traces*) at three different baseline membrane potentials (−45 mV *light blue*, −40 mV *purple*, and −35mV red). Current contribution to I_KVTOT_ from K_V_1.5 (indicated by asterisks) and K_V_2.1 in male and female myocytes at a baseline membrane potential of −45 mV (**C**), −40 mV (**D**), and - 35 mV (**E**).

Having explored the regulation of graded membrane potential by the activation of I_KVTOT_ to counterbalance the nonselective cations currents (I_NSC_), we next explored the effects of steady membrane depolarization in the *in silico* vascular smooth muscle cell myocyte model on I_Ca_ in male and female myocytes. We predicted I_Ca_ in our male and female simulations at steady-state membrane depolarization after simulation for 500 seconds. We observed that as the membrane depolarizes from −55 to −35 mV, I_Ca_ in male myocytes increased from 0 to 1.0 pA while in female myocytes I_Ca_ increased from 0 to 1.5 pA as shown in **Figure 7A**, suggesting that I_Ca_ is larger in female compared to those of male myocytes. We recorded the predicted [Ca^2+^]_i_ and observed that I_Ca_ led to a higher calcium influx in female compared to male simulations as shown in **Figure 7B**. To illustrate in detail, we show in **Figure 7C-D**, time traces of *in silico* predictions of membrane voltage at −40 mV (*top panel*), I_Ca_ (*middle panel*), and [Ca^2+^]_i_ (*lower panel*) corresponding to the male and female data points indicated by black and blue arrows respectively shown in **Figure 7A-B**. In the male case (**Figure 7C**), at a steady membrane potential of −40 mV, L-type calcium Ca_V_1.2 channels produced a current of 0.5 pA. However, in female simulations (**Figure 7D**), we observed that at a steady membrane potential of −40 mV, L-type calcium Ca_V_1.2 channels produced a current of 0.65 pA. We calculated that at −40 mV, two Ca_V_1.2 channels are needed to sustain 0.5 pA of current in male myocytes while three Ca_V_1.2 channels are needed to sustain 0.65 pA of current in female myocytes. Although the sex-specific differences in male and female simulations at −40 mV are small, a 15 nM difference in the overall response of [Ca^2+^]_i_ can have a profound effect on the constriction state of the myocytes. The predictions from the Hernandez-Hernandez model provide a comprehensive picture of physiological conditions and support the idea that a small number of Ca_V_1.2 channels supply the steady Ca^2+^ influx needed to support a maintained constricted state in small arteries and arterioles^53,69^. The differences between males and females are notable in the context of observations indicating varied sex-based responses to antihypertensive agents that target the Ca^2+^ handling system in vascular smooth muscle cells.

**Figure 7.**
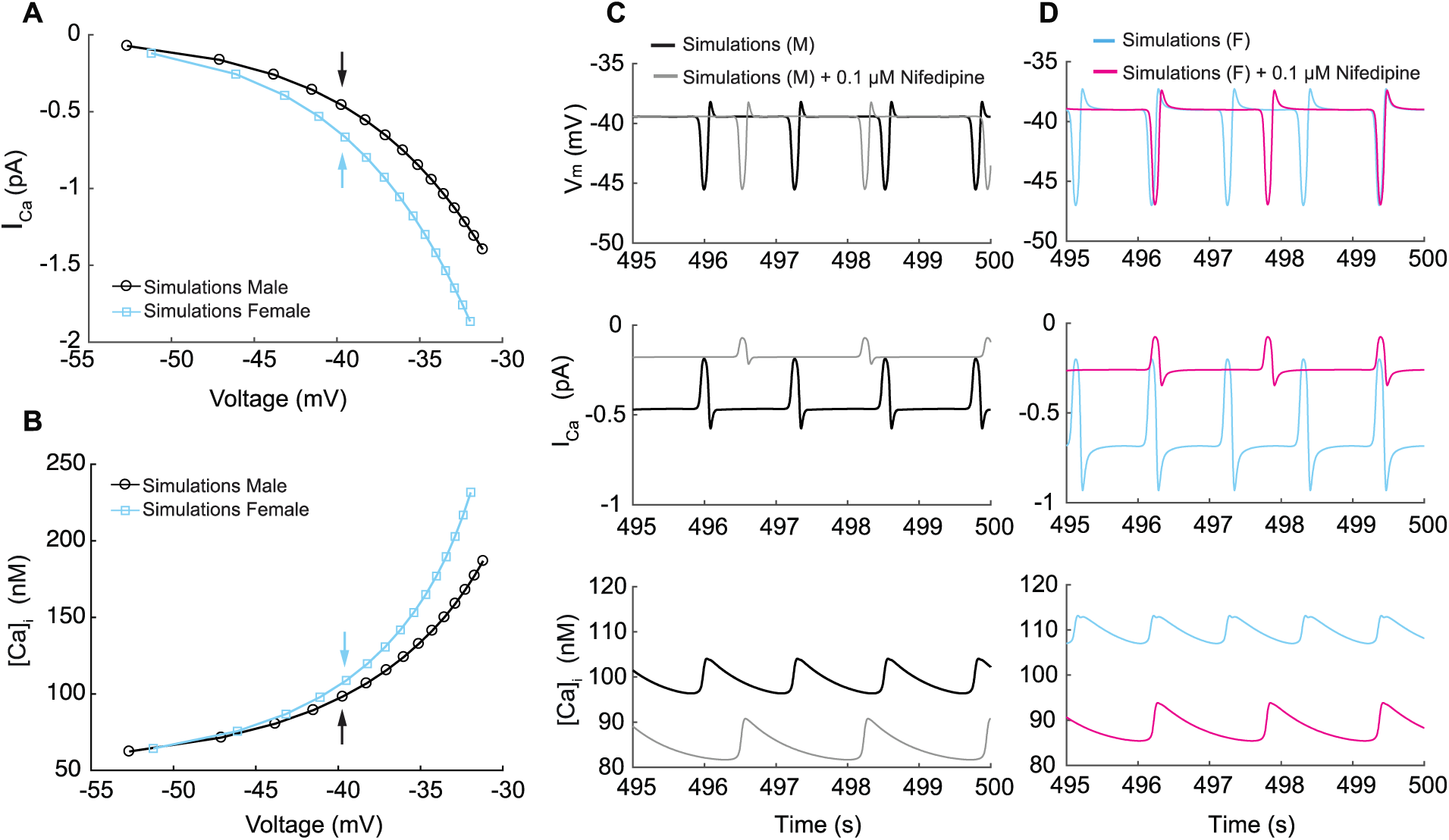
Simulated L-type calcium currents (I_Ca_) and calcium influx in male and female vascular smooth muscle cells. **A**) Male and female whole-cell I_Ca_ membrane potential relationship. (**B**) Male and female intracellular calcium concentration in the cytosolic compartment at indicated membrane potential. **C**) Time course of membrane potential in male vascular smooth muscle cells before (black) and after (gray) simulated nifedipine application (*top panel*). Corresponding time course of L-type calcium current I_Ca_ before (black) and after (gray) simulated nifedipine application (*middle panel*) and intracellular calcium [Ca^2+^]_i_ concentration before (black) and after (gray) simulated nifedipine application (*lower panel*). **D**) Time course of membrane potential in female vascular smooth muscle cells before (blue) and after (pink) simulated nifedipine application (*top panel*). Corresponding time course of L-type calcium current I_Ca_ before (blue) and after (pink) simulated nifedipine application (*middle panel*) and intracellular calcium [Ca^2+^]_i_ concentration before (blue) and after (pink) simulated nifedipine application (*lower panel*).

Next, we simulated the effects of calcium channel blocker nifedipine on I_Ca_ at a steady membrane potential of −40 mV in male and female simulations. Briefly, previous studies^70^ have shown that at the therapeutic dose of nifedipine (i.e., about 0.1 μM) L-type Cav1.2 channel currents are reduced by about 60-70%. Accordingly, we decreased I_Ca_ in our mathematical simulations by the same extent. In **Figure 7C-D**, we show the predicted male (gray) and female (pink) time course of membrane voltage at −40 mV (*top panel*), I_Ca_ (*middle panel*), and [Ca^2+^]_i_ (*lower panel*). First, we observed that in both male and females 0.1 μM nifedipine modifies the frequency of oscillation in the membrane potential, by causing a reduction in oscillation frequency. Second, both male and female simulations (middle panels) show that 0.1 μM nifedipine caused a reduction of I_Ca_ to levels that are very similar in male and female myocytes following treatment. Consequently, the reduction of I_Ca_ causes both male and female simulations to reach a very similar baseline [Ca^2+^]_i_ of about 85 nM (lower panels). As a result, simulations provide evidence supporting the idea that Ca_V_1.2 channels are the predominant regulators of intracellular [Ca^2+^] entry in the physiological range from −40 mV to −20 mV. Importantly, these predictions also suggest that clinically relevant concentrations of nifedipine cause larger overall reductions in Ca^2+^ influx in female than in male arterial myocytes.

Thus far, we have shown the development and application of models of vascular smooth muscle myocytes incorporating measured sex-specific differences in currents from male and female isolated cells. Given that hypertension is essentially a consequence of the spatial organization and function of smooth muscle cells^71,72^, we next expanded our study to include a one-dimensional (1D) tissue representation of electrotonically coupled tissue by connecting arterial myocytes in series.

A well-known phenomenon in excitable systems is that electrotonic coupling between cells results in the minimization of individual cellular differences, thereby producing a smoothing effect across the tissue^73–75^. We simulated 400 female or 400 male vascular smooth muscle myocytes and set the gap junctional conductivity to zero to uncouple the simulated cells. As expected, the uncoupled cells in both male and female cases demonstrated the characteristic behavior of arterial myocytes, exhibiting spontaneous hyperpolarization. Of the 400 cells, we show the simulated representative traces of Cell 1, Cell 50, and Cell 100 for female (**Figure 8A**) and male (**Figure 8B**).

**Figure 8.**
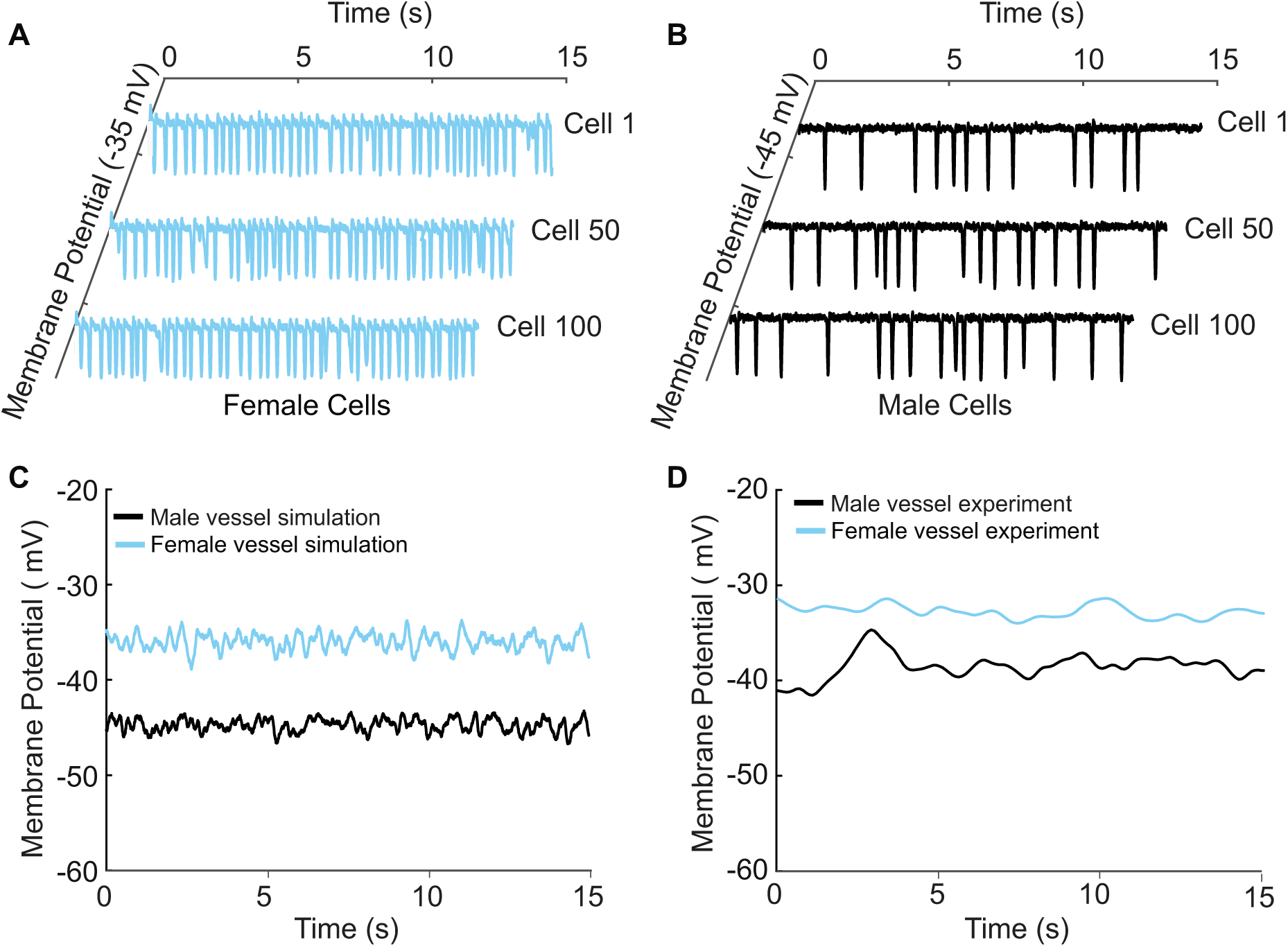
A one-dimensional tissue model representation of vascular smooth muscle cells connected in series. **A**) Uncoupled female vessel simulation showing cell 1, cell 50, and cell 100 at a baseline membrane potential of −35 mV. **B**) Uncoupled male vessel simulations showing cell 1, cell 50, and cell 100 at a baseline membrane potential of −45 mV. **C**) Composite female (blue trace) and male (black trace) membrane potential of 400 coupled smooth muscle cells connected with gap junctional resistance of 71.4 Ωcm^2^ in a one-dimensional tissue representation. (**D**) Sharp-electrode records of the membrane potential of smooth muscle in pressurized (80-mmHg) female and male arteries from O’Dwyer *et al.*, 2020.

Next, we modeled 400 cells but with electrotonic coupling by setting the simulated gap junctional resistance to 71.4 Ωcm^2 76^. In this case, we observed that the spontaneous hyperpolarizations, previously observed in the uncoupled cells, diminished when cells were coupled. The overall smoothing effect observed in **Figure 8C** is attributed to the electrotonic coupling and consequential influence of neighboring cells. The electrical response is consistent across the spatial domain for both male (**Figure 8C**; *black trace*) and female (**Figure 8C**-*blue trace*) one-dimensional tissue representations. Notably, the model predicts a more depolarized female membrane potential in the one-dimensional tissue representations consistent with experimental measurements as shown in **Figure 8D**.

Having developed an idealized model of a vessel, we set out to validate the model predictions of variable [Ca^2+^]_i_ between males and females by comparing the computed calcium signaling in vascular smooth muscle with experimental recordings O’Dwyer *et al.*^20^. Given that membrane potential predominantly governs calcium influx in vascular smooth muscle^14^, we varied the conductance of the nonselective cation currents (I_NSCC_) in our simulations. Tuning of I_NSCC_ was performed to replicate the effects of pressure-induced membrane depolarization, which results in activation of the voltage-gated L-type Ca^2+^ channels and increases [Ca^2+^]_i_.

Our simulations (lines) are well validated by experimental recordings (symbols) in **Figure 9A**. A distinctive feature from the model prediction, which was validated by experimental recordings is the observation that female (**Figure 9A**, blue trace and symbols) vessels accommodate more [Ca^2+^]_i_ compared to male (**Figure 9A**, black trace and symbols) vessels. Intriguingly, the mechanism of different [Ca^2+^]_i_ in male and female vessels was revealed in single-cell simulations, which showed attributable sex-based differences in L-type Ca^2+^ currents.

**Figure 9.**
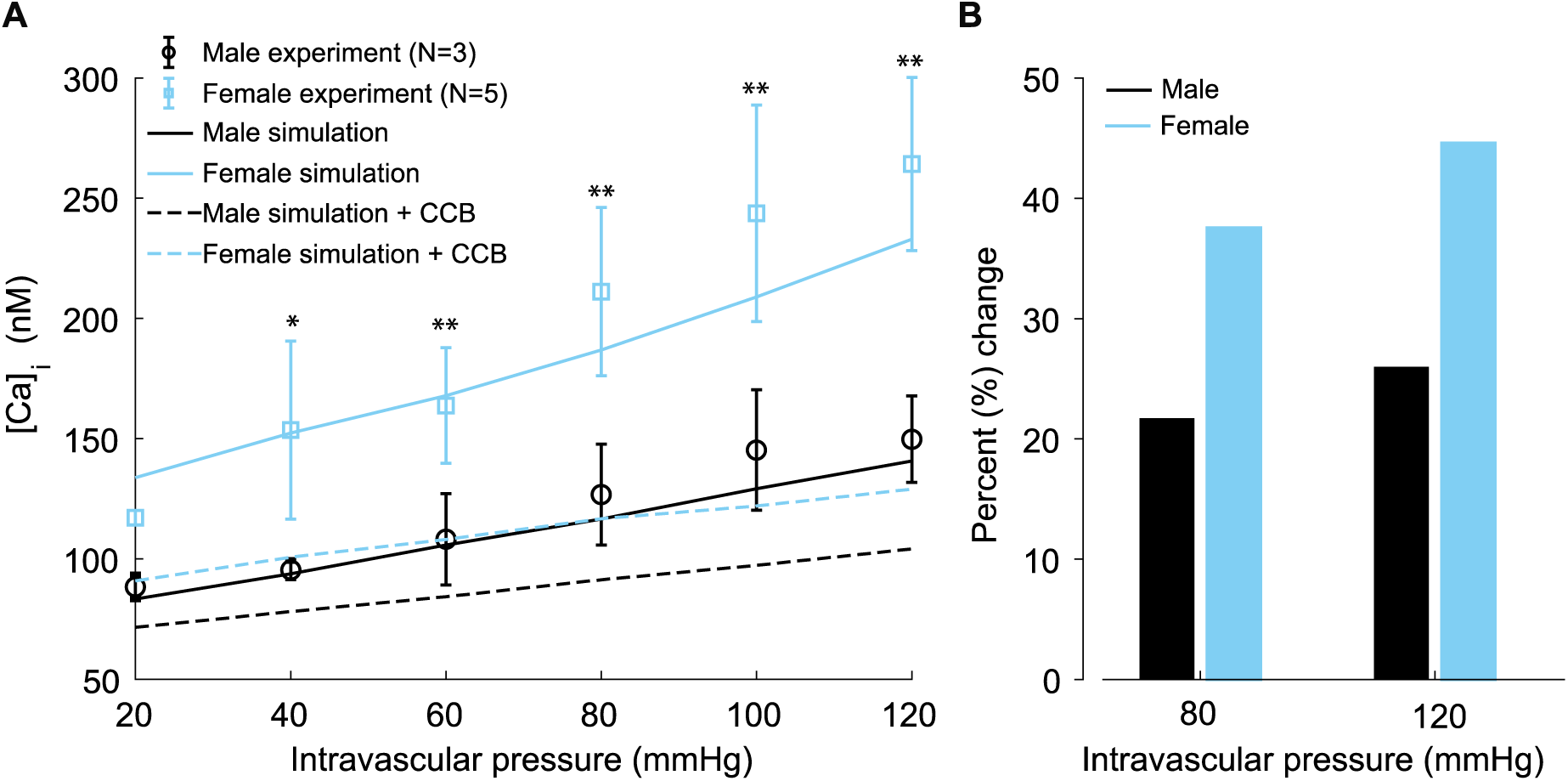
Experimentally measured and modeled intracellular calcium [Ca]_i_ in male and female vessels and response to clinically used L-type Ca^2+^ channel blocker. **A**) Intracellular calcium [Ca]_i_ in female (blue symbols) and male (black symbols) arteries at intravascular pressures ranging from 20 to 120 mmHg. Simulations showing [Ca]_i_ in the idealized female and male vessels are shown with blue and black solid lines, respectively. Simulated [Ca]_i_ after the application of clinically used L-type Ca^2+^ channel blocker nifedipine is shown with dashed lines for male (black) and female (blue). **B**) Comparison of the percentage change of [Ca]_i_ in male (black) and female (blue) after the application L-type Ca^2+^ channel blocker nifedipine at 80 mmHg and 120 mmHg. *P < 0.05, **P < 0.01.

Finally, in our simulations, we computed the effects of [Ca^2+^]_i_ after the application of clinically relevant calcium channel blocker nifedipine. We observed a substantial reduction of [Ca^2+^]_i_ in both male (**Figure 9A**, *dashed black line*) and female (**Figure 9A**, *dashed blue line*). Significant differences were found in the physiological range of intravascular pressure from 40 to 120 mmHg. In the summary data (**Figure 9B**), we quantified the relative change of [Ca^2+^]_i_ in male (*black*) and female (*blue*) after the application of 0.1 mM L-type Ca^2+^ channel blocker nifedipine at 80 mmHg and 120 mmHg. Our results show that nifedipine, when applied to male vessels, decreases [Ca^2+^]_i_ by 22% and 25% at 80 mmHg and 120 mmHg, respectively. However, the same dose of nifedipine when applied to female vessels decreases [Ca^2+^]_i_ by 38% and 45% at 80 mmHg and 120 mmHg. The results suggest that female arterial smooth muscle is more sensitive to clinically used Ca^2+^ channel blockers than male smooth muscle.

## Discussion

Here, we describe the development, validation, and application of an *in-silico* model to simulate and understand the mechanisms of electrical activity and Ca^2+^ dynamics in a single mesenteric vascular smooth muscle cell. The Hernandez-Hernandez model is the first model to incorporate sex-specific differences in voltage-gated K_V_2.1 and Ca_V_1.2 channels and predicts sex-specific differences in membrane potential and Ca^2+^ signaling regulation in the smooth muscle of both sexes from systemic arteries. In the pursuit of stratifying sex-specific responses to antihypertensive drugs, we expanded our exploration to encompass a one-dimensional (1D) tissue representation. Such an approach allowed us to simulate and predict the impact of Ca^2+^ channel blockers within a mesenteric vessel. Notably, the computational framework can be expanded to forecast the impact of antihypertensives and other perturbations from single-cell to tissue-level simulations.

To specifically investigate the impact of sex-specific differences measured from ion channel experiments and their impact on membrane potential and [Ca^2+^]_i_, we focused on the isolated myocyte in the absence of complex signaling pathways. We first explored the effects of Ca_V_1.2 and K_V_2.1 channels on membrane potential as experimental data suggest key sex-specific differences in channel expression and kinetics. Notably, the peak of the current-voltage (I-V) relationship of L-type Ca_V_1.2 current is 40% smaller in male compared to female myocytes (**Figure 2D**).

Similarly, the peak current-voltage (I-V) relationship of the voltage-gated K_V_2.1 current (I_Kv2.1_) is 70% smaller in male compared to female myocytes at +40 mV (**Figure 3C**). O’Dwyer and coauthors^20^ showed sex-dependent expression of K_V_2.1 in the plasma membrane, where male arterial myocytes have a total of about 75,000 channels compared to 183,000 channels in female myocytes. Notably, less than 0.01% of channels are conducting in male and female myocytes. In the computational model, we found that to reproduce the experimentally measured amplitude of the K_V_2.1 I-V curve (**Figure 3C**), a maximum of ∼44 male K_V_2.1 channels was sufficient to reproduce the peak current (68.8 pA at 40 mV). In contrast, ∼143 channels were predicted to be needed in female myocytes to reproduce the experimentally measured peak current (226.42 pA at +40 mV) of the K_V_2.1 I-V relationship. Modeling and simulation led to the prediction that in male arterial myocytes, I_KvTOT_ is largely dictated by K_V_1.5 channels. In contrast, in female arterial myocytes, K_V_2.1 channels dominate I_KvTOT_ **(Figure 3F)**.

An important aspect of the Hernandez-Hernandez model is that it includes Ca^2+^-mediated signaling between RyRs in the junctional SR and BK_Ca_ channel clusters in the nearby sarcolemma membrane. This section of the model is similar to the one included in the Karlin model^5^ with some modifications. The Karlin model described how subcellular junctional spaces influence membrane potential and [Ca^2+^]_i_ in response to intravascular pressure, vasoconstrictors, and vasodilators. In this study, we reduced the complexity of the model representation of subcellular Ca^2+^ signaling spaces to include just three compartments: the cytosol, SR, and the SR-sarcolemma junction. Our model represents on average, the behavior of a single junctional SR unit that is functional in a cell at a time. The model uses a deterministic approach but mimics the process of production of Ca^2+^ sparks that activate BK_Ca_ channel clusters^29^. We represented the activity of the RyRs in the junctional domain deterministically in the model so that Ca^2+^ spark-BK_Ca_ currents occur at a frequency of about 1 Hz at −40 mV in a space equivalent to 1% of the total cell surface area of the plasma membrane^22^.

Based on experimental observations, the Hernandez-Hernandez computational model employs three key assumptions: First, Ca^2+^ sparks in the junctional domain are initiated by activation of RyRs, where RyR gating opening probability is correlated with SR load. Second, Ca^2+^sparks lead to a [Ca^2+^]_Jun_ increase between 10-20 mM to match the amplitude measured in experiments (**Figure 4A**)^54,58^. Third, activation of BK_Ca_ channels and the resultant current amplitude derives from the experimentally observed spontaneous outward currents (STOCs) in both amplitude and morphology.

Notably, model simulations revealed important mechanisms that may underlie experimental observations in measurements of membrane potential (**Figure 5A**). The model predicts that the mechanism of intrinsic oscillatory behavior in the vascular myocytes results from a delicate balance of currents. Activation of non-selective cation currents (I_NSC_) likely causes membrane depolarization, but the delayed rectifier currents (I_KvTOT_) oppose them, resulting in membrane potential baseline in the physiological range of −45 to −20 mV. Interestingly, the voltage-gated L-type Ca_V_1.2 currents activation threshold sits within this range at ∼-45 mV. Therefore, small increases in I_NSC_ can overwhelm I_KvTOT_ below −20 mV and result in sufficient depolarization to bring the membrane potential to the threshold for activation of I_Ca_. It is important to note that I_KvTOT_ increases sharply upon depolarization from −45 to −20 mV, resulting in tight control of membrane potential and prevention of large transient depolarization resulting from I_NSC_. Activation of L-type Ca^2+^ channels upon depolarization and subsequent Ca^2+^ release within the small volume junction then activates the BK_Ca_ channels, which results in hyperpolarization. Hyperpolarization reignites the oscillatory cascade as an intrinsic resetting mechanism. Since vascular myocytes are subject to substantial noise from the stochastic opening of ion channels in the plasma membrane, and fluctuations in the local junctional domain components, such as the SR load, RyR opening, and BK_Ca_ channel activity, we included noise in the simulation. To simulate the physiological noise in the vascular smooth muscle cell (**Figure 5B**), we added Gaussian noise to the dV/dt equations and [Ca]_SR_.

Female mesenteric artery myocytes are more depolarized than male myocytes at physiological intravascular pressures^20^. Our model suggests that female myocytes are more depolarized than male myocytes due to larger non-selective cation currents in female compared to male myocytes, most likely due to the activation of Na^+^-permeable TRP channels. To our knowledge, the only TRP channels found to regulate the membrane potential of mesenteric artery smooth muscle are TRPP1 and TRPP2^11,12^. Future work will have to determine if TRPC6^8^ and TRPM4^9,10^, which have been shown to mediate the myogenic response of cerebral artery smooth muscle, and/or other non-selective cation channels also depolarize mesenteric artery smooth muscle^77^.

The Hernandez-Hernandez model predicts that very few channels (based on total current amplitude) are likely to control the baseline fluctuations in membrane potential in the physiological range of −60 to −20 mV. The intrinsic oscillatory properties of the vascular myocyte operating in the low voltage regime under conditions of high resistance membrane are similar to other types of oscillatory electrical cells including cardiac pacemaker cells.

As shown in (**Figure 6**), the model predicts that, at −40 mV, the amplitude of steady-state K_V_2.1 currents is about 0.8 pA in male and 3.3 pA in female arterial myocytes, indicating that the contribution of K_V_2.1 and K_V_1.5 channels to membrane potential is different in males and females. At −30 mV, it is 2.34 pA and 9.2 pA in male and female myocytes respectively. Assuming a single channel current at −40 and −30 mV of 0.7 pA, we calculated that, on average, in male myocytes a single channel is open at −40 mV and 3 channels are open at any particular time at −30 mV. In female myocytes, 6 channels are predicted to be open at −40 mV, while 13 are predicted to be active at −30 mV.

The Hernandez-Hernandez model also allowed us to calculate the number of Ca_V_1.2 channels needed to sustain the steady-state concentration of [Ca^2+^]_i_ in the physiological range from −60 to −20 mV (**Figure 7**). The model predicts that at −40 mV in mouse male myocytes, 2 channels were required to generate 0.5 pA of steady-state Ca_V_1.2 current. On the other hand, we found that in female myocytes, 3 channels were sufficient to generate 0.65 pA of Ca_V_1.2 current. These data are consistent with the work of Rubart et al.^69^, which suggested that steady-state Ca^2+^ currents at −40 mV were likely produced by the opening of 2 Ca_V_1.2 channels in rat cerebral artery smooth muscle cells.

The observation that a very small number of the conducting K_V_2.1 and Ca_V_1.2 channels are involved in the regulation of membrane potential and Ca^2+^ influx in male and female arterial myocytes at physiological membrane potentials is important for several reasons. *First*, the analysis suggests that small differences in the number of K_V_2.1 and Ca_V_1.2 channels can translate into large, functionally important differences in membrane potential and [Ca^2+^]_i_ and hence affect and control myogenic tone under physiological and pathological conditions. *Second*, the small number of K_V_2.1 and Ca_V_1.2 channels gating between −40 and −30 mV likely makes smooth muscle cells more susceptible to stochastic fluctuations in the number and open probabilities of these channels than in cells where a large number of channels regulate membrane excitability and Ca^2+^ influx (e.g., adult ventricular myocyte^78^). This, at least in part, likely contributes to Ca^2+^ signaling heterogeneity in vascular smooth muscle.

Hypertension fundamentally manifests through the spatial organization of cellular components, particularly evident in the context of the tunica media, the middle layer of vessels is predominantly constituted of smooth muscle cells which play a pivotal role in vessel contraction and relaxation^71,72^. Such intricate biological machinery is imperative in orchestrating the regulation of blood flow and blood pressure. Our approach began with a process of distillation, aiming to shed light on cellular mechanisms within isolated vascular myocytes from small systemic vessels and arterioles, which control blood pressure, of both male and female mice.

Earlier research has confirmed that in mesenteric arteries, the pathogenesis leading to hypertension is largely determined by the downregulation of K_V_2.1^36^ and/or K_V_1.5^67,79^ and a concurrent increase in the activity of Ca_V_1.2^80^ channels. Building upon this knowledge, we broadened our study to encompass a one-dimensional (1D) tissue model of electrotonically linked tissue, achieved by connecting arterial myocytes in series. The 1D cable model has anatomical relevance because the structure of third and fourth-order mesenteric arteries have a singular layer of vascular myocytes encircling the lumen in a cylindrical arrangement. The cable structure is analogous to an “unrolled” or lateral arrangement of the vessel. Such an approach allowed a conceptual framework to bridge the gap between understanding the combined effects of membrane potential and [Ca^2+^]_i_ in isolated cells and in the wider context of small vessels.

For instance, previous studies have proposed that gap junctions enable vessels to function in a way that is analogous to a large capacitor^57,81^. The gap junctions actively filter and transform single-cell electrical activity into sustained responses across the tissue^81^. Recent studies add to this understanding by demonstrating that Connexin 37 (Cx37), a component of these gap junctions, seems to be expressed in the mesenteric arteries^82^. In our simulations, we showed (**Figure 8A-B**) that indeed uncoupled cells exhibit a spontaneous oscillatory behavior which studies have confirmed is not an artifact due to isolation from the vessel but rather an intrinsic behavior required to sustain electrical signals. When the cells are connected (**Figure 8C**) the spontaneous hyperpolarization previously observed in the uncoupled cells diminished, the effect is attributed to the electrotonic coupling and consequential influence of neighboring cells. In addition, in our simulations, we found that it is required to have stochastic fluctuations to allow the system to average the membrane potential behavior that dictates the amount of [Ca^2+^]_i_ in the vessels.

Regarding, Ca_V_1.2 channels, simulations forecast the clinically relevant concentrations (0.1 μM) at which common Ca^2+^ channel blockers (e.g., nifedipine) effectively block Ca_V_1.2 channels in both male and female smooth muscle (**Figure 9**). Our simulations in isolated arterial myocytes and in the one-dimensional (1D) tissue model suggest heightened sensitivity to calcium channel blockers in the female compared to male.

The model predictions are aligned with documented sex-specific differences in antihypertensive drug responses^83,84^. Previous studies, notably by Kloner et al., have illustrated this point quantitatively, highlighting a more pronounced diastolic BP response in women (91.4%) compared to men (83%) when treated with dihydropyridine-type channel blockers, such as amlodipine. Importantly, this distinction persisted even after adjusting for confounding factors such as baseline BP, age, weight, and dosage per kilogram^84^. Another interesting observation from Kajiwara et al. emphasizes that vasodilation-related adverse symptoms occur more frequently in younger women (<50 years) compared to their male counterparts, again suggesting a heightened sensitivity to dihydropyridine-type calcium channel blockers^85^.

## Limitations

The model presented here describes the necessary and sufficient ion channels, pumps, and transporters to describe the electrical activity and Ca^2+^ signaling of an isolated mesenteric smooth muscle cell in the absence of complex signaling pathways. Such an approach enabled us to perform a component dissection to analyze the sex-specific differences observed in the fundamental electrophysiology of male and female myocytes. However, it is well known that vascular smooth muscle cells are subject to a plethora of stimuli from endothelial cells, neurotransmitters, endocrine, and paracrine signals^5^. The next phase of the project includes an expansion of the model to incorporate receptor mediated signaling pathways that are essential for blood pressure control.

Excitation-contraction coupling refers to an electrical stimulus that drives the release of calcium from the sarcoplasmic reticulum and results in the physical translocation of fibers that underlies muscle contraction. In the present model, we did not explicitly consider the mechanical description of muscle contraction. Nevertheless, we can imply contractile effects by tracking membrane potential and the elevation of [Ca^2+^]_i_ as a proxy.

To conclude, we developed and present the Hernandez-Hernandez model of male and female isolated mesenteric vascular myocytes. An additional limitation of our study is the reliance on predominantly murine data. Although mouse arteries do present numerous parallels with human arteries—including analogous intravascular pressure-myogenic tone relationships, resting membrane potentials, and the expression of typical ionic channels like Ca_V_1.2, BKCa channels, and RyRs^86–88^. Future research should assess the direct applicability and implication of our findings in human subjects.

## Conclusions

The Hernandez-Hernandez model of the isolated mesenteric vascular myocyte was informed and validated with experimental data from male and female vascular myocytes. We then used the model to reveal sex-specific mechanisms of K_V_2.1 and Ca_V_1.2 channels in controlling membrane potential and Ca^2+^ dynamics. In doing so, we predicted that very few channels are needed to contribute to and sustain the oscillatory behavior of the membrane potential and calcium signaling. We expanded our computational framework to include a one-dimensional (1D) tissue representation, providing a basis for simulating the effect of drug effects within a vessel. The model predictions suggested differences in the response of male and female myocytes to drugs and the underlying mechanisms for those differences. These predictions may constitute the first step towards better hypertensive therapy for males and females.

## MATERIALS AND METHODS

### Section 1. Experimental

#### 1.1 Animals

This study was performed in strict accordance with the recommendations in the Guide for the Care and Use of Laboratory Animals of the National Institutes of Health. All of the animals were handled according to approved institutional animal care and use committee (IACUC) protocols of the University of California Davis. IACUC protocol number is 22503. 8- to 12-week-old male and female mice C57BL/6J (The Jackson Laboratory, Sacramento, CA) were used in this study. Animals were housed under standard light-dark cycles and allowed to feed and drink ad libitum. Animals were euthanized with a single lethal dose of sodium pentobarbital (250 mg/kg) intraperitoneally. All experiments were conducted in accordance with the University of California Institutional Animal Care and Use Committee guidelines.

8- to 12-week-old male and female mice C57BL/6J (The Jackson Laboratory, Sacramento, CA) were used in this study. Animals were housed under standard light-dark cycles and allowed to feed and drink ad libitum. Animals were euthanized with a single lethal dose of sodium pentobarbital (250 mg/kg) intraperitoneally. All experiments were conducted in accordance with the University of California Institutional Animal Care and Use Committee guidelines.

#### 1.2 Isolation of arterial myocytes from systemic resistance arterioles

Third and fourth-order mesenteric arteries were carefully cleaned of surrounding adipose and connective tissues, dissected, and held in ice-cold dissecting solution (Mg^2+^-PSS; 5 mM KCl, 140 mM NaCl, 2mM MgCl_2_, 10 mM glucose, and 10 mM HEPES adjusted to pH 7.4 with NaOH). Arteries were first placed in dissecting solution supplemented with 1.23 mg/ml papain (Worthington Biochemical, Lakewood, NJ) and 1 mg/ml DTT for 14 minutes at 37°C. This was followed by a second 5-minute incubation in dissecting solution supplemented with 1.6 mg/ml collagenase H (Sigma-Aldrich, St. Louis, MO), 0.5 mg/ml elastase (Worthington Biochemical, Lakewood, NJ), and 1 mg/ml trypsin inhibitor from *Glycine max* (Sigma-Aldrich, St. Louis, MO) at 37°C. Arteries were rinsed three times with dissection solution and single cells were obtained by gentle trituration with a wide-bore glass pipette. Myocytes were maintained at 4°C in dissecting solution until used.

#### 1.3 Patch-clamp electrophysiology

All electrophysiological recordings were acquired at room temperature (22–25°C) with an Axopatch 200B amplifier and Digidata 1440 digitizer (Molecular Devices, Sunnyvale, CA). Borosilicate patch pipettes were pulled and polished to resistances of 3-6 MΩ for all experiments using a micropipette puller (model P-97, Sutter Instruments, Novato, CA).

Voltage-gated Ca^2+^ currents (I_Ca_) were measured using conventional whole-cell voltage-clamp sampled at a frequency of 50 kHz and low-pass filtered at 2 kHz. Arterial myocytes were continuously perfused with 115 mM NaCl,10 mM TEA-Cl, 0.5 mM MgCl_2_, 5.5 mM glucose, 5 mM CsCl, 20 mM CaCl_2_, and 10 mM HEPES, adjusted to pH 7.4 with CsOH. Micropipettes were filled with an internal solution containing 20 mM CsCl, 87 mM Aspartic acid, 1 mM MgCl_2_, 10 mM HEPES, 5 mM MgATP, and 10 mM EGTA adjusted to pH 7.2 using CsOH. Current-voltage relationships were obtained by exposing cells to a series of 300 ms depolarizing pulses from a holding potential of −70 mV to test potentials ranging from −70 to +60 mV. A voltage error of 9.4 mV due to the liquid junction potential of the recording solutions was corrected offline. Voltage dependence of Ca^2+^ channel activation (G/G_max_) was obtained from the resultant currents by converting them to conductance via the equation G = I_Ca_/(test potential – reversal potential of I_Ca_); normalized G/G_max_ was plotted as a function of test potential. Time constants of activation and inactivation of I_Ca_ were fitted with a single exponential function.

I_Kv_ recordings were performed in the whole-cell configuration with myocytes exposed to an external solution containing 130 mM NaCl, 5 mM KCl, 3 mM MgCl_2_, 10 mM Glucose, and 10 mM HEPES adjusted to 7.4 using NaOH. The internal pipette solution constituted of 87 mM K-Aspartate, 20 mM KCl, 1 mM CaCl_2_, 1 mM MgCl_2_, 5 mM MgATP, 10 mM EGTA, and 10 mM HEPES adjusted to 7.2 by KOH. A resultant liquid junction potential of 12.7 mV from these solutions was corrected offline. To obtain current-voltage relationships cells were subjected to a series of 500 ms test pulses increasing from −70 to +70 mV. To isolate the different K^+^ channels attributed to composite I_K_, cells were first bathed in external I_K_ solution, subsequently exposed to 100 nM Iberiotoxin (Alomone, Jerusalem, Israel) to eliminate any BK_Ca_ channel activity and finally immersed in an external solution containing both 100 nM Iberiotoxin and 100 nM Stromatoxin (Alomone, Jerusalem, Israel) to block both BK_Ca_ and K_V_2.1 activity. Ionic current was converted to conductance via the equation G = I(V-E_K_). E_K_ was calculated to be −78 mV. Activation time constants for K_V_2.1 currents were obtained by fitting the rising phase of these currents with a single exponential function.

BK_ca_-mediated spontaneous transient outward currents (STOCs) and membrane potential were recorded using the perforated whole-cell configuration. To measure both, myocytes were continuously exposed to a bath solution consisting of 130 mM NaCl, 5 mM KCl, 2 mM CaCl_2_, 1 mM MgCl_2_, 10 mM glucose, and 10 mM HEPES, pH adjusted to 7.4 with NaOH. Pipettes were filled with an internal solution containing 110 mM K-aspartate, 30 mM KCl, 10 mM NaCl, 1 mM MgCl_2_, 0.5 mM EGTA, and 10 mM HEPES adjusted to a pH of 7.3 with KOH. The internal solution was supplemented with 250 µg/ml amphotericin B (Sigma, St. Louis, MO). STOCs were measured in the voltage-clamp mode and were analyzed with the threshold detection algorithm in Clampfit 10 (Axon Instruments, Inc). Membrane potential was measured using the current clamp mode.

STOCs were recorded using the perforated whole-cell configuration. The composition of the external bath solution consisted of 134 mM NaCl, 6 mM KCl, 1 mM MgCl_2_,2 mM CaCl_2_,10 mM glucose, and 10 mM HEPES adjusted to a pH of 7.4 using NaOH. Pipettes were filled with an internal solution of 110 mM K-aspartate, 10 mM NaCL, 30 mM KCl, 1 mM MgCl_2_, 160 μg/ml amphotericin B, and 10 mM HEPES using NaOH to adjust to pH to 7.2. Myocytes were sustained at a holding potential of −70 mV before being exposed to a 400 ms ramp protocol from −140 to +60 mV. A voltage error of 12.8 mV resulting from the liquid junction potential was corrected for offline. K_ir_ channels were blocked using 100 µM Ba^2+^.

#### 1.4 Statistics

Data are expressed as mean ± SEM. All data sets were tested for normality. Normally distributed data were analyzed using T-tests or one-way analysis of variance (ANOVA). ANOVA analyses were followed by multiple comparison tests (i.e. Tuckey). P<0.05 was considered statistically significant.

### Section 2. Computational modeling and simulation

#### 2.1 Cell size and structure

The mean capacitance of the cells was experimentally calculated to be 16±3 pF based on all the male and female WT mesenteric C57BL/6J cells utilized in the experiments (N=45). Assuming the cells are roughly cylindrical in shape, the expected radius should be 2.485 *μm* and a length of 100 *μm* leading to a surface area of 1.6×10^-5^ cm^2^ and a total volume of approximately 1.94×10^-12^ liters. The cell capacitance of excitable membranes is assumed to be 1.0×10^-6^ F/cm^2^, with the calculated surface area the estimated total cell capacitance is C_m_ = 16 pF.

Because the total cell volume is roughly 2×10^-12^ liters, it is assumed that 50% of the total cell volume is occupied by organelles. There are three main compartments in vascular myocytes important to the regulation of membrane potential and calcium signaling: the cytosol, sarcoplasmic reticulum (SR), and specialized junctional domains formed by the SR and the plasma membrane. The cytosol occupies approximately 50% of total cell volume (V_cyt_ = 1.0×10^-12^ L). The sarcoplasmic reticulum occupies approximately 5% of cell cytosolic volume (V_SR_ = 5.0×10^-14^ L), and the junctional domain volume is approximately 1% of the cytosol volume (V_Jun_ = 0.5×10^-14^ L)^22,37–39^.

#### 2.2 Model development

The male and female *in silico* models are single whole-cell models based on the electrophysiology of isolated mesenteric vascular smooth muscle myocytes. A schematic of the proposed model is shown in **Figure 1**. The membrane electrophysiology can be described by the differential equation:

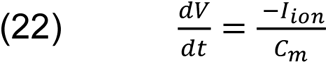

Where V is voltage, t is time, C_m_ is membrane capacitance I_ion_ is the sum of transmembrane currents. The contribution of each transmembrane current to the total transmembrane ionic current can be described by the following equation:

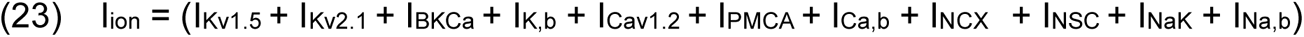

The eleven transmembrane currents are generated by ion channels, pumps, and transporters. Currents from ion channels include the voltage-gated L-type calcium current (I_Ca_), the nonselective cation current (I_NSC_), voltage-gated potassium currents (I_Kv1.5_ and I_Kv2.1_), and the large-conductance Ca^2+^-sensitive potassium current (I_BKCa_). Additionally, there are three background or leak currents (I_K,b_, I_Ca,b_, and I_Na,b_). Currents from pumps and transporters include the sodium-potassium pump current (I_NaK_), plasma membrane Ca-ATPase transport current (I_PMCA_), and sodium-calcium exchanger current (I_NCX_).

Cytosolic concentrations of sodium and potassium as a function of time are determined by considering the sum of their respective fluxes into the cytosol.

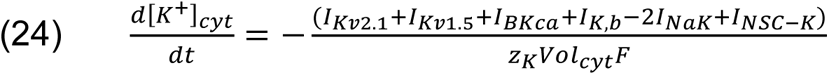

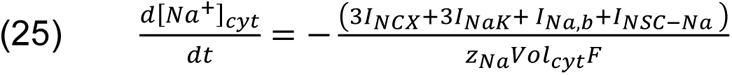

Where F is the Faraday’s constant, Vol_cyt_ is the cytoplasmic volume and *Z*_*K*_and *Z*_*Na*_ are the valence of potassium and sodium ions, respectively.

The calcium dynamics is compartmentalized into three distinct regions: cytosol [Ca^2+^]_i_, the sarcoplasmic reticulum [Ca^2+^]_SR_, and the junctional region [Ca^2+^]_jun_. The cytosol includes a calcium buffer, which we assume can be described as a first-order dynamics process.

*Cytosolic calcium region (*[Ca^2+^]_i_): calcium concentration in this region varies between 100-300 nM^13^ and is mainly influenced by the following fluxes: transmembrane pumps and transporters, the sarcoplasmic reticulum Ca-ATPase (J_SERCA_), diffusion from the junctional domain region (J_Jun-Cyt_) and the calcium buffer calmodulin (BUF_CAM_).

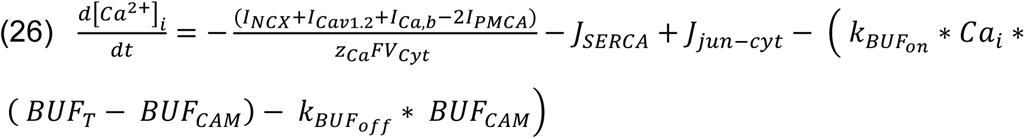

*Sarcoplasmic reticulum region ([Ca^2+^]_SR_):* calcium concentration in this region varies between 100-150 μM^40^ and it is mainly influenced by the sarcoplasmic reticulum Ca-ATPase (J_SERCA_) and the flux from the ryanodine receptors (J_RyR_).

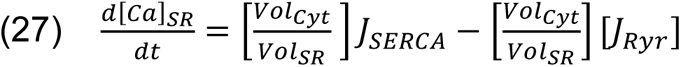

*Junctional region ([Ca^2+^]_jun_):* calcium concentration in this region varies between 10-100 μM^37,38^ and is mainly influenced by the flux from the ryanodine receptors (J_RyR_), the diffusion from the junctional region to the cytoplasm (J_Jun-Cyt_)

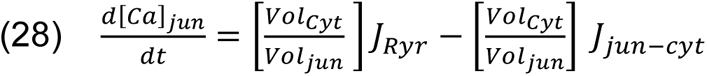

In the model, the flux of J_SERCA_ was adapted from the Luo-Rudy II model^41^ and the flux of J_RyR_ was adapted from previous models of ryanodine receptors activation, originally introduced in the field of cardiac electrophysiology^42–44^.

#### 2.3 Parameter optimization and reformulation of the gating ion channel models

The ionic current models of I_Ca_, I_Kv2.1_, and I_Kv1.5_ were optimized using the approach employed by Kernik *et al*.^45^. Here, the open probability P_o_ of each voltage-dependent gating variable “n” was defined by opening- and closing-rate voltage-dependent functions α_n_ and β_n_ respectively, and were modeled by simple exponentials of the form:

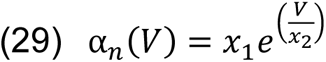

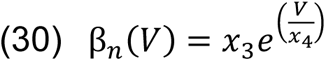

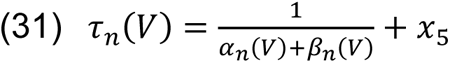

The steady-state availability remains the same as the classical Hodgkin-Huxley formulations and the time constant values follow a modified version formulation by accommodating an extra parameter x_5_ in equation 31. (x_1_, x_2_, x_3_, x_4_, x_5_) are parameters to be optimized using experimental data. We used the parameter optimization employed by Kernik *et al.*^45^, which minimizes the error between model and experimental data using the Nelder–Mead minimization of the error function. Random small perturbations (<10%) were applied to find local minima, to improve data fit. The parameter fit with the minimal error function value after 1000 to 10000 perturbations was used as the optimal model fit to the data.

#### 2.4 Cellular simulations with noise

The simulations encapsulate the cumulative effect of stochastic ion channel activity on cell voltage dynamics through the fluctuating current term, ξ(t), into the membrane potential (dV/dt) equation^46^, as shown in equation 32. Here it is assumed ξ(t) is only a function of time and it is implemented as Gaussian white noise^47^.

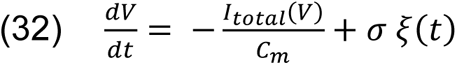

We use the Euler-Maruyama numerical method for updating equation 33 and 34 as follows.

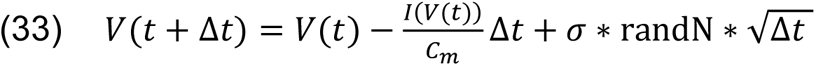

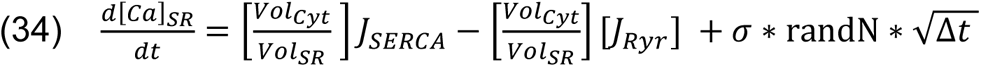

Where randN is a random number from a normal distribution (N(0,1)) with mean 0 and variance 1. Δ*t* is the time step and *σ* is the “diffusion coefficient”, which represents the amplitude of the noise. The numerical method for updating the voltage was forward Euler.

#### 2.5 One-dimensional simulations

The idealized one-dimensional representation of a vessel was developed by connecting 400 Hernandez-Hernandez model cells in series via simulated resistances to represent gap junctions. For each cell in the cable, the Hernandez-Hernandez model was used to compute ionic currents and concentration changes. The temporal transmembrane fluxes of the Hernandez-Hernandez model are related to the spatial or current flow by a finite difference approximation of the cable equation^48–50^

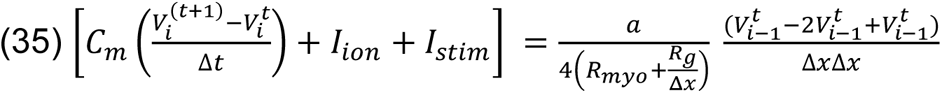

where I_ion_ represents the individual membrane ionic current densities (pA/pF) of the Hernandez-Hernandez model, I_stim_ is the stimulus current density (pA/pF) set to zero in our simulation, *a* is the radius of the fiber (5 μm), C_m_ is the membrane capacity (pA/pF), V_it_ is membrane potential at segment i and time t, Δx is the discretization element (100 μm = 0.01 cm). Where R_myo_ is the myoplasmic resistance (R_myo_=150 Ωcm) and R_g_ is the gap junction resistance (R_g_=71.4 Ωcm^2^).

#### 2.6 Sensitivity Analysis

The baseline models in male and female vascular smooth muscle cells were analyzed through a parameter sensitivity assessment using multivariable linear regression, following the methodology introduced by Sobie^51^. The scope of the sensitivity analysis encompassed variations in the maximal conductance and maximal ion transport rates of the transmembrane currents, including I_Kv1.5_, I_Kv2.1_, I_BKCa_, I_K,b_, I_Cav1.2_, I_PMCA_, I_Ca,b_, I_NCX_, I_NSC_, I_NaK_, and I_Na,b_. All other parameters, notably those defining model kinetics, remained constant at the values established by the foundational model. Scaling factors were randomly selected from a log-normal distribution characterized by a median value of 1 and a standard deviation of 0.1^45^.

#### 2.7 Simulation protocols

Code for simulations and analysis was written in C++ and MATLAB 2018a. The single vascular smooth muscle code was run on an Apple Mac Pro machine with two 2.7 GHz 12-Core Intel Xeon processors and an HP ProLiant DL585 G7 server with a 2.7 GHz 28-core AMD Opteron processor. Vessel simulations were implemented in C++ and parallelized using OpenMP. The C++ code was compiled with the Intel ICC compiler, version 18.0.3. Numerical results were visualized using MATLAB R2018a by The MathWorks, Inc. All codes and detailed model equations are available on GitHub (https://github.com/ClancyLabUCD/sex-specific-responses-to-calcium-channel-blockers-in-mesenteric-vascular-smooth-muscle)

## Acknowledgments

This work was supported by National Institutes of Health Common Fund Grant OT2OD026580 (to C.E.C., L.F.S.), National Heart, Lung, and Blood Institute grants R01HL128537 (C.E.C, L.F.S.,), R01HL152681 (to C.E.C., L.F.S.). UC Davis Department of Physiology and Membrane Biology Research Partnership Fund (to C.E.C.) as well as UC Davis T32 Predoctoral and Postdoctoral Training in Basic and Translational Cardiovascular Medicine fellowship supported in part by NHLBI Institutional Training Grant T32HL086350 (to C.M., G.H.H.)

